# Saliva-Based Molecular Testing for SARS-CoV-2 that Bypasses RNA Extraction

**DOI:** 10.1101/2020.06.18.159434

**Authors:** Diana Rose E. Ranoa, Robin L. Holland, Fadi G. Alnaji, Kelsie J. Green, Leyi Wang, Christopher B. Brooke, Martin D. Burke, Timothy M. Fan, Paul J. Hergenrother

## Abstract

Convenient, repeatable, large-scale molecular testing for SARS-CoV-2 would be a key weapon to help control the COVID-19 pandemic. Unfortunately, standard SARS-CoV-2 testing protocols are invasive and rely on numerous items that can be subject to supply chain bottlenecks, and as such are not suitable for frequent repeat testing. Specifically, personal protective equipment (PPE), nasopharyngeal (NP) swabs, the associated viral transport media (VTM), and kits for RNA isolation and purification have all been in short supply at various times during the COVID-19 pandemic. Moreover, SARS-CoV-2 is spread through droplets and aerosols transmitted through person-to-person contact, and thus saliva may be a relevant medium for diagnosing SARS-CoV-2 infection status. Here we describe a saliva-based testing method that bypasses the need for RNA isolation/purification. In experiments with inactivated SARS-CoV-2 virus spiked into saliva, this method has a limit of detection of 500-1000 viral particles per mL, rivalling the standard NP swab method, and initial studies also show excellent performance with 100 clinical samples. This saliva-based process is operationally simple, utilizes readily available materials, and can be easily implemented by existing testing sites, thus allowing for high-throughput, rapid, and repeat testing of large populations.

**Graphical Abstract:** 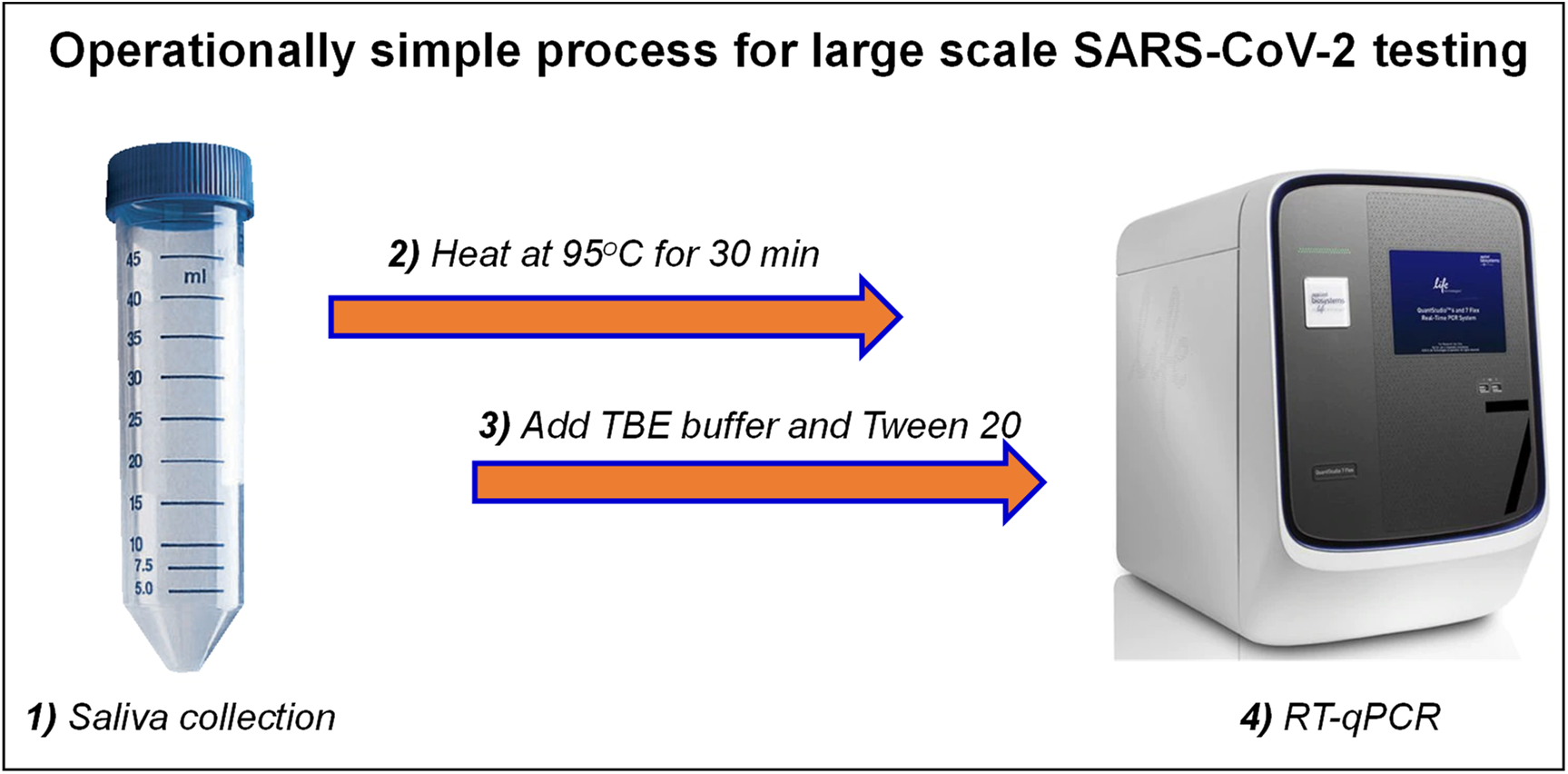

## Background

The slow roll-out and inconsistent availability of diagnostic testing for SARS-CoV-2 has hobbled efforts to control the COVID-19 pandemic in many countries. Testing protocols based on the use of nasopharyngeal (NP) swabs as the collection agent, placed in a tube containing viral transport media (VTM), followed by RNA isolation/purification and subsequent analysis by RT-qPCR is currently the most common method (**Figure 1A**).^1,2^ While some variant of this process has been implemented worldwide, there are multiple challenges with this workflow. Sample collection using NP swabs requires healthcare workers wearing personal protective equipment (PPE) to collect samples, the swabs can be uncomfortable for the patients during collection, and the swabs and the associated VTM have been in short supply at many times and in most locations. In addition, RNA isolation/purification is another significant bottleneck, both in the time and labor required for this process, and in the availability of the equipment and reagents. All of these components also add to the cost of the testing process.

**Figure 1.**
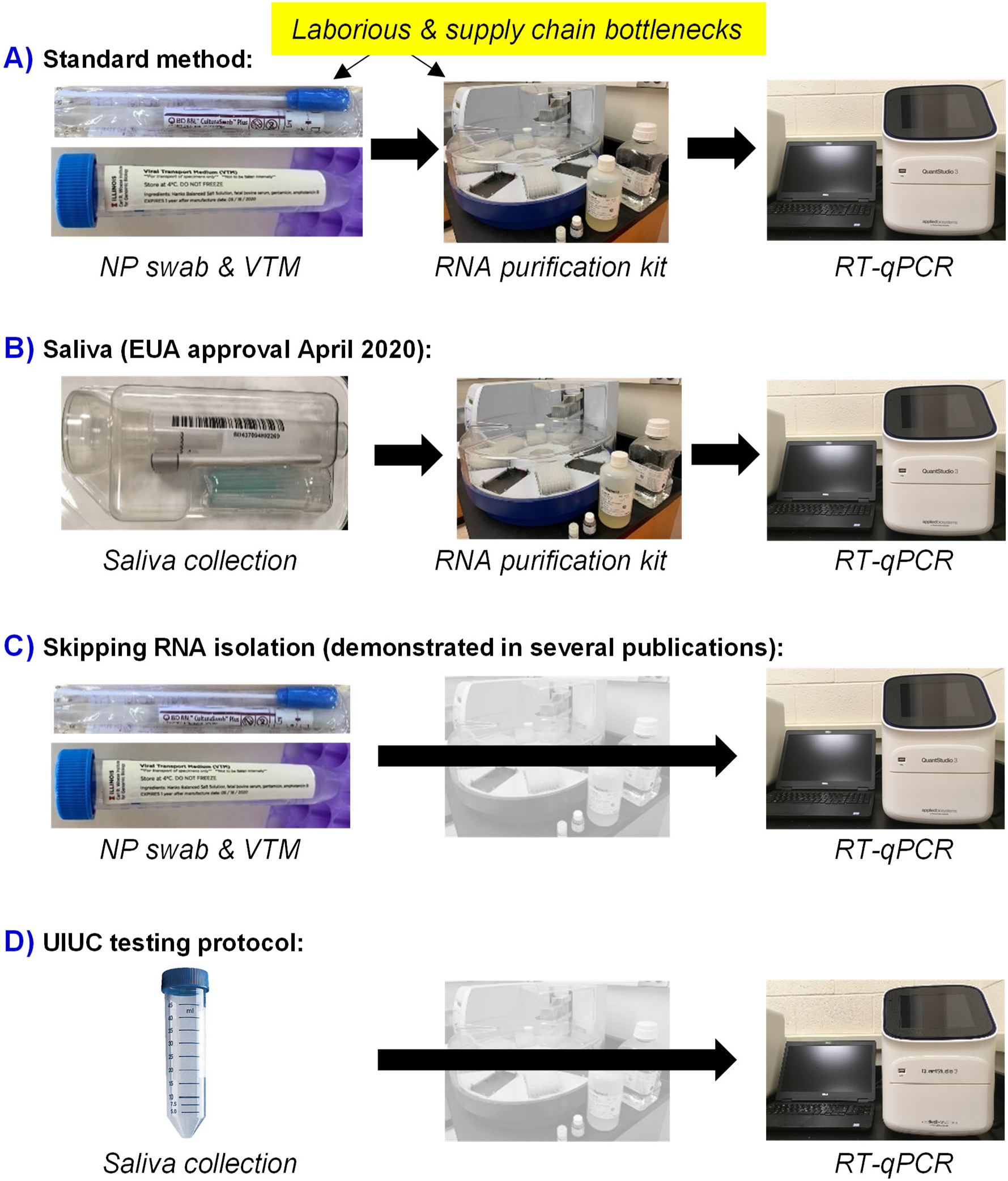
Schematic of SARS-CoV-2 testing. **A)** The current, widely-utilized diagnostic process involves nasopharyngeal (NP) swabs and viral transport media (VTM), followed by RNA extraction and isolation, with RT-qPCR analysis of the samples. NP swabs, VTM, and RNA purification kits have been in short supply at various times. **B)** In April of 2020, saliva was emergency use authorized (EUA) as a diagnostic sample, using RNA extraction and isolation, followed by RT-qPCR. **C)** Other groups have reported direct testing of NP swabs in VTM by RT-qPCR. **D)** The University of Illinois at Urbana-Champaign (UIUC) protocol involves saliva collection in standard 50 mL conical tubes, heating (95°C for 30 min), followed by addition of buffer and analysis by RT-qPCR.

There is emerging consensus that widespread, frequently repeated testing is necessary for a safer return to activities that are important for society. Given the data suggesting that SARS-CoV-2 can be spread by pre-symptomatic/asymptomatic carriers,^3–6^ localized outbreaks could be dramatically reduced or prevented if individuals shedding SARS-CoV-2 could be readily identified and isolated. For example, imagine a testing bubble placed over a group that desires face-to-face interaction – employees of a company, members of a sports team, extended family networks, etc. If all members of the group could be tested for SARS-CoV-2, then isolated, then tested again after an appropriate time increment (likely ~4-5 days, in line with the incubation period for SARS-CoV-2^7,8^), two negative tests would provide confidence for a safer return to activities. Of course, in practice there are challenges with total self-isolation and avoidance of others outside the testing bubble, but the above scenario represents one promising path forward, allowing positive cases to be identified and contained, and reducing the probability that pre-symptomatic/asymptomatic virus shedders unknowingly transmit SARS-CoV-2 to others. Unfortunately, as the size of a group grows larger, widespread and frequent testing for SARS-CoV-2 using the standard testing protocol depicted in **Figure 1A** becomes impractical. For example, it would be untenable to repeatedly test all members of a university in a short time period using this process, and thus we were motivated to develop a streamlined, cost-effective, SARS-CoV-2 testing platform that can be realistically scaled to test thousands of individuals a day.

When considering various sample collection possibilities, saliva is attractive due to the known detection of SARS-CoV-2 through oral shedding, and the potential for rapid and easy self-collection,^9–11^ thus minimizing the need for direct healthcare provider-patient contact and consequent conservation of PPE. In addition, a number of recent reports have detailed the detection of SARS-CoV-2 in saliva through the workflow in **Figure 1B**, including a report showing higher viral loads in saliva when compared to matched NP swabs from the same patients.^12^ Importantly, saliva (expelled in aerosols and droplets) may be a significant factor in person-to-person transmission of SARS-CoV-2,^10^ and it has been suggested that NP swab tests remain positive long after patients are infectious (potentially due to detection of inactive virus or remnants of viral RNA in the NP cavity),^13^ whereas SARS-CoV-2 viral loads in saliva are highest during the first week of infection, when a person is most infectious. These data suggest that viral loads in saliva may be a good reflection of the transmission potential of patients infected SARS-CoV-2.^13–15^

While we are unaware of direct SARS-CoV-2 detection from saliva that bypasses RNA isolation/purification, there are several reports of detection from swab/VTM that bypasses RNA isolation/purification (**Figure 1C**).^16–23^ With the ultimate goal of providing convenient, scalable, and costeffective molecular diagnostic testing for >10,000 individuals per day using a single COVID-19 testing center, here we report the discovery of a sensitive saliva-based detection method for SARS-CoV-2 that bypasses RNA isolation/purification (**Figure 1D**). This SARS-CoV-2 testing process and workflow is convenient, simple, rapid, and inexpensive, and can be readily adopted by any testing facility currently using RT-qPCR.

## Results

### Development of a direct saliva-to-RT-qPCR process for detection of SARS-CoV-2

While SARS-CoV-2 has been identified in the nasopharynx, collecting NP samples is neither trivial nor innocuous, and for repeat testing to track disease progression within a given patient this method may prove unreliable, due to inconsistencies in repeated sampling and potential formation of scar tissue, altogether resulting in possible false-negatives.^24^ Compounding these anatomic limitations, the procedure for NP sample collection is invasive, further reducing patient compliance for repeated and serial sampling. Saliva may serve as an important mediator in transmitting SARS-CoV-2 between individuals via droplets and aerosols,^25–27^ and thus viral loads in saliva may serve as a highly relevant correlate of transmission potential. However, saliva is comprised of constituents that may hinder virus detection by RT-qPCR, such as degradative enzymes. As such, we sought to identify conditions that could take advantage of the many positives of saliva while overcoming potential limit of detection challenges with this collection medium. For the optimization phase of this work we utilized two versions of inactivated SARS-CoV-2, one inactivated through gamma(γ)-irradiation (5×10^6^ RADs) and one inactivated through heat (65°C, 30 min). For the detection of SARS-CoV-2, we utilized the commercially available TaqPath RT-PCR COVID-19 kit, developed and marketed by Thermo Fisher Scientific. This multiplex RT-qPCR kit targets the ORF1ab (replication), N-gene (nucleocapsid), and S-gene (spike) of SARS-CoV-2. To reduce cost and extend reagent usage, we performed RT-qPCR reactions at half the suggested reaction mix volume.^28^

#### Heat treatment

Up-front heating of freshly collected saliva samples is attractive as a simple method to inactivate the virus without having to open the collection vessel. Indeed, heat treatment is often used to inactivate saliva patient samples,^29,30^ thus conferring added biosafety by decreasing the likelihood of viral transmission via sample handling by personnel. Common conditions for SARS-CoV-2 inactivation are heating at 56-60°C for 30-60 min,^30,31^ although other temperature and times have been examined.^30^ Using intact, γ-irradiated SARS-CoV-2 spiked into fresh human saliva (that was confirmed to be SARS-CoV-2 negative), we observed dramatic time- and temperature-dependent improvement in SARS-CoV-2 detection by direct RT-qPCR, without the use of RNA extraction. When incubated at ambient temperature (no heat treatment), no SARS-CoV-2 genes were detectable (**Figure 2**). As temperature and incubation time were increased, substantial improvement in virus detection was observed, with 100% identification of all SARS-CoV-2 genes, in all replicate samples, being detected following a 30 min incubation at 95°C. Importantly, a short heating time (5 minutes) at 95°C (as has been examined by others^29,32^) does not allow for sensitive detection; the 30 minute duration is essential, as it is likely that this extended heating inactivates components of saliva that inhibit RT-qPCR. Thus, proper heating of patient samples allows for virus detection without the need for RNA extraction, with the added benefit of inactivating the samples, thus substantially reducing biohazard risks.

**Figure 2.**
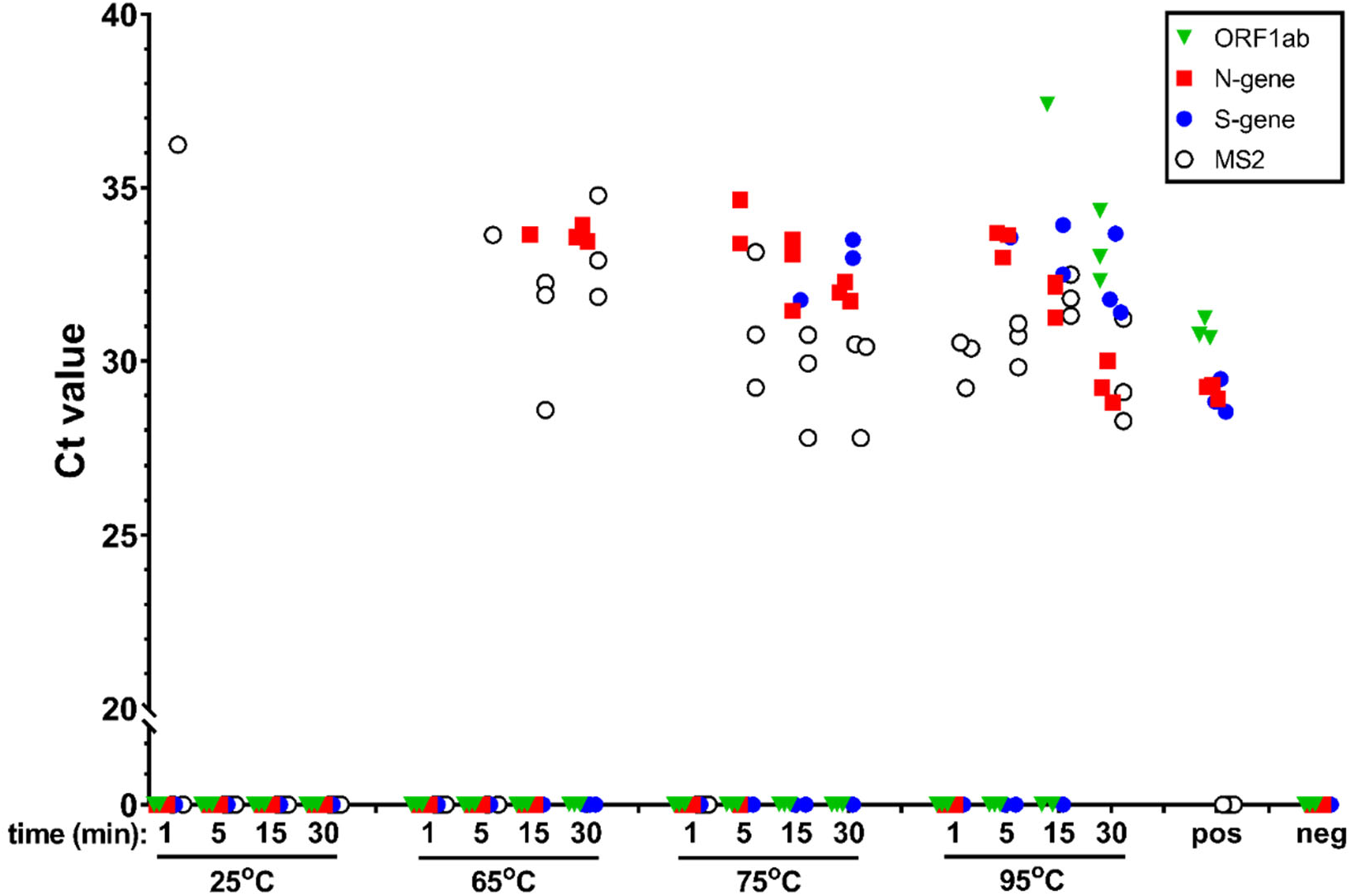
The effect of heat on SARS-CoV-2 detection. γ-irradiated SARS-CoV-2 (from BEI, used at 1.0×10^4^ viral copies/mL) was spiked into fresh human saliva (SARS-CoV-2 negative). Samples diluted 1:1 with 2X Tris-borate-EDTA (TBE) buffer (0.5 mL in 50 mL conical tubes) were incubated at 25°C (ambient temperature), or in a hot water bath at 65°C, 75°C, or 95°C, for 1, 5, 15, or 30 min. All saliva samples were spiked with purified MS2 bacteriophage (1:40 MS2:sample) as an internal control. Virus-spiked saliva samples, a positive control (pos; SARS-CoV-2 positive control, 5.0×10^3^ copies/mL, no MS2) and a negative control (neg; water, no MS2) were directly analyzed by RT-qPCR, in triplicate, for SARS-CoV-2 ORF1ab (green triangle), N-gene (red square), and S-gene (blue circle), and MS2 (open circle). Undetermined Ct values are plotted at 0.

#### Saliva collection buffer

We next sought to evaluate saliva collection buffers as a means to enhance viral RNA stability, but also to increase uniformity between saliva samples and to decrease sample viscosity. In conjunction with RNA isolation/purification, other groups have utilized protocols whereby saliva was provided by a patient and soon thereafter combined with the collection buffer; reported collection buffers include Phosphate Buffered Saline (PBS),^33^ DNA/RNA Shield,^34^ and Tris-EDTA (TE).^35^ Using intact, γ-irradiated SARS-CoV-2 spiked into fresh human saliva, which was then heat treated at 95°C for 30 min, we observed outstanding virus detection when saliva samples were combined with either Tris-Borate-EDTA (TBE) or TE buffer (**Figure 3A**). Comparable Ct values were observed between TBE and TE buffer, but TE yielded greater variability between individual gene replicates, whereas TBE buffer yielded highly clustered data. In stark contrast, combining saliva with PBS or two commercially available buffers (DNA/RNA Shield, SDNA-1000), completely abrogated viral detection, including the MS2 bacteriophage internal control, indicating that these buffers directly interfere with the RT-qPCR reaction itself. TBE, TE, and PBS were further titrated with different concentrations of SARS-CoV-2, where similar trends were observed, namely, greater replicate variability with TE buffer, and no virus detection with PBS (**Supporting Figure 1**). Thus, when saliva samples are combined with TBE buffer to a final working concentration of 1X, SARS-CoV-2 is detectable in saliva without RNA extraction; TE buffer is also suitable but more variability is observed. These findings further suggest that while PBS and commercially available buffers may be appropriate for samples that are processed via RNA extraction, these agents are incompatible with direct saliva-to-RT-qPCR.

**Figure 3.**
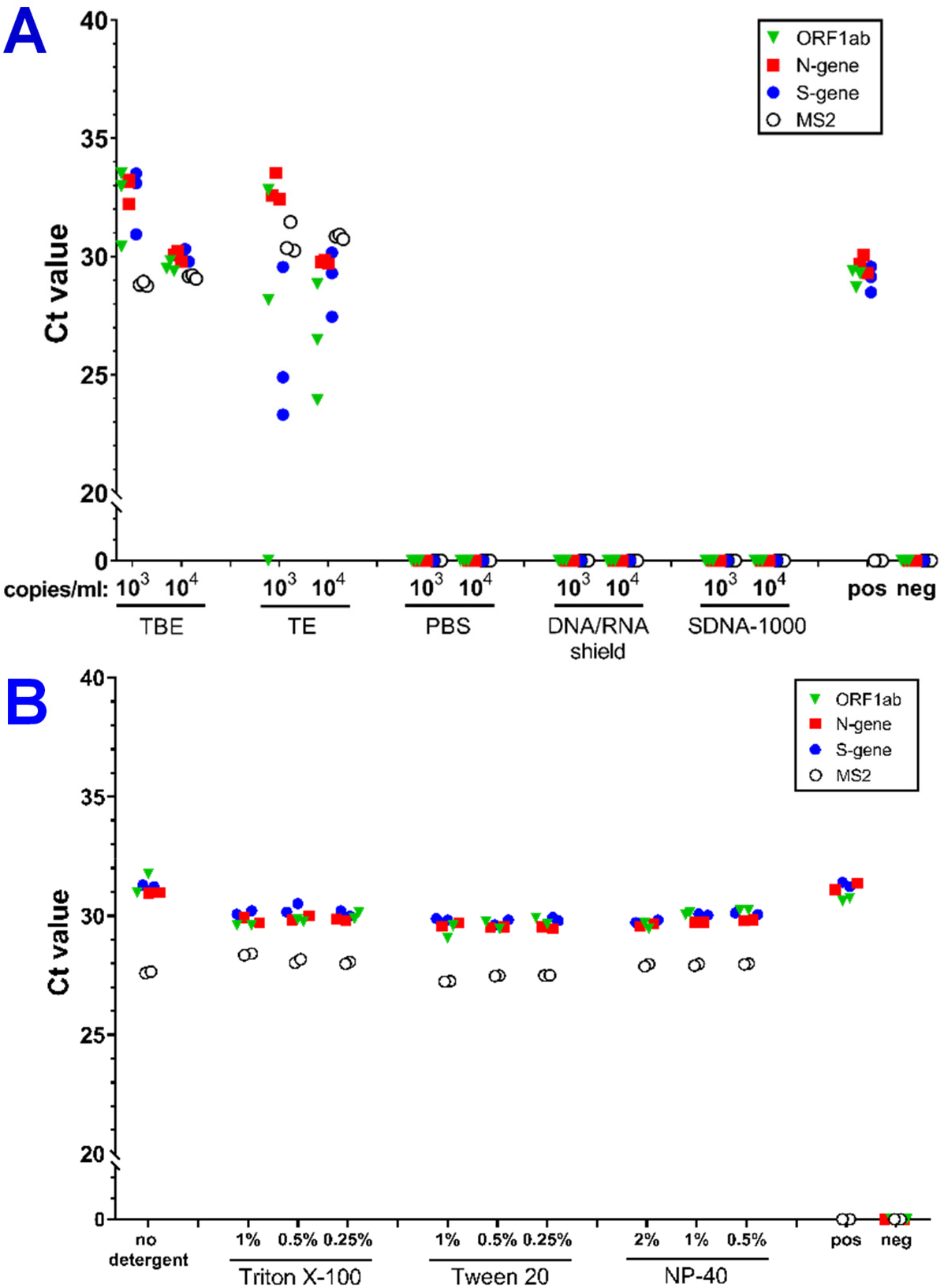
(**A**) The effect of collection buffer on SARS-CoV-2 detection. γ-irradiated SARS-CoV-2 (from BEI, at 1.0×10^3^ or 1.0×10^4^ viral copies/mL) was spiked into fresh human saliva (SARS-CoV-2 negative) and combined at a 1:1 ratio with Tris-Borate-EDTA (TBE), Tris-EDTA (TE), Phosphate Buffered Saline (PBS), DNA/RNA shield (Zymo Research), or SDNA-1000 (Spectrum Solutions) such that the final concentration of each buffer was 1X. Samples (0.5 mL in 50 mL conical tubes) were incubated in a hot water bath at 95°C for 30 min. (**B**) Detergent optimization. γ-irradiated SARS-CoV-2 (1.0×10^4^ viral copies/mL) was spiked into fresh human saliva (SARS-CoV-2 negative) and combined 1:1 with TBE buffer at a final working concentration of 1X. Samples were treated with detergents (Triton X-100, 1%, 0.5%, 0.25%; Tween 20, 1%, 0.5%, 0.25%; NP-40, 2%, 1%, 0.5%) after heating at 95°C for 30 min. All saliva samples were spiked with purified MS2 bacteriophage (1:40 MS2:sample) as an internal control. Virus-spiked saliva samples, a positive control (pos; SARS-CoV-2 positive control, 5.0×10^3^ copies/mL, no MS2) and a negative control (neg; water, no MS2) were directly analyzed by RT-qPCR, in triplicate, for SARS-CoV-2 ORF1ab (green triangle), N-gene (red square), and S-gene (blue circle), and MS2 (open circle). Undetermined Ct values are plotted at 0.

#### Sample additives

In addition to saliva collection buffers, various additives have been explored for their ability to enhance SARS-CoV-2 detection.^1,36–38^ Therefore, detergents, including Triton X-100, Tween 20, and NP-40 (**Figure 3B**), as well as various RNA stabilizing agents, including RNase inhibitor, carrier RNA, glycogen, TCEP, proteinase K, bovine serum albumin (BSA), RNAlater, and PBS-DTT (**Supporting Figure 2**) were examined. Notably, modest improvements in viral detection were observed with all detergents tested (~2 Ct, **Figure 3B**) and with addition of carrier RNA, RNase inhibitor, and BSA (**Supporting Figure 2**), These additives slightly improve virus detection, without interfering with RT-qPCR; in addition, if clinical saliva specimens are especially viscous, addition of detergent may improve ease of sample handling. However, inclusion of detergents prior to heat treatment inhibited viral detection, emphasizing the importance of adding detergents after heat treatment, if they are to be included (**Supporting Figure 3**). Of the detergents tested, Tween 20 was chosen for incorporation into the standard sample processing protocol, given its ease of handling and cost. When samples were treated with Tween 20 and TBE (alone or in combination, either before or after heating) the ideal workflow for virus detection, as defined by the lowest Ct values with the greatest clustering of individual replicates, was TBE buffer before heating, and Tween 20 after heating (**Supporting Figure 3**). However, it is important to note that comparable results were obtained when TBE was added after heating (**Supporting Figure 3**), suggesting flexibility in when TBE buffer can be included during sample processing. Altogether, the safest and most streamlined protocol would be: collection of saliva samples, heat at 95°C for 30 min, add TBE buffer and Tween 20, followed by RT-qPCR.

#### Limit of detection

Using the optimized protocol of addition of TBE (or TE) buffer at a 1:1 ratio with saliva, followed by heat treatment at 95°C for 30 min and addition of Tween 20 to a final concentration of 0.5%, the limit of detection (LOD) was determined. Other reports have suggested that SARS-CoV-2 is shed into saliva at a remarkably wide range from 10,000-10,000,000,000 copies/mL.^12,26^ While the LOD of SARS-CoV-2 approved diagnostic methods can vary considerably (500-80,000 viral copies/mL^39^) and are not always reported, the best LOD values for SARS-CoV-2 using RNA extraction protocols appear to be approximately 1000 copies/mL.^28^ Similarly, a LOD of 5610 copies/mL was found for SARS-CoV-2 detection in saliva using RNA purification.^12^ To determine the LOD for this new direct protocol (saliva➔RT-qPCR), a side-by-side comparison was conducted of intact, γ-irradiated SARS-CoV-2 spiked into fresh human saliva compared to a process that includes RNA isolation/purification. As shown in **Figure 4**, comparable LOD measurements were observed, with LOD of ~500 viral copies/mL for both the direct process with addition of Tween 20 and TBE buffer, and the process using RNA purification. Similar results were observed with heat-inactivated SARS-CoV-2, whereby the LOD was measured to be 5000 viral copies/mL for both RNA extraction of saliva samples and direct saliva-to-RT-qPCR, with greater detection if the virus was directly analyzed in water (**Supporting Figure 4**).

**Figure 4.**
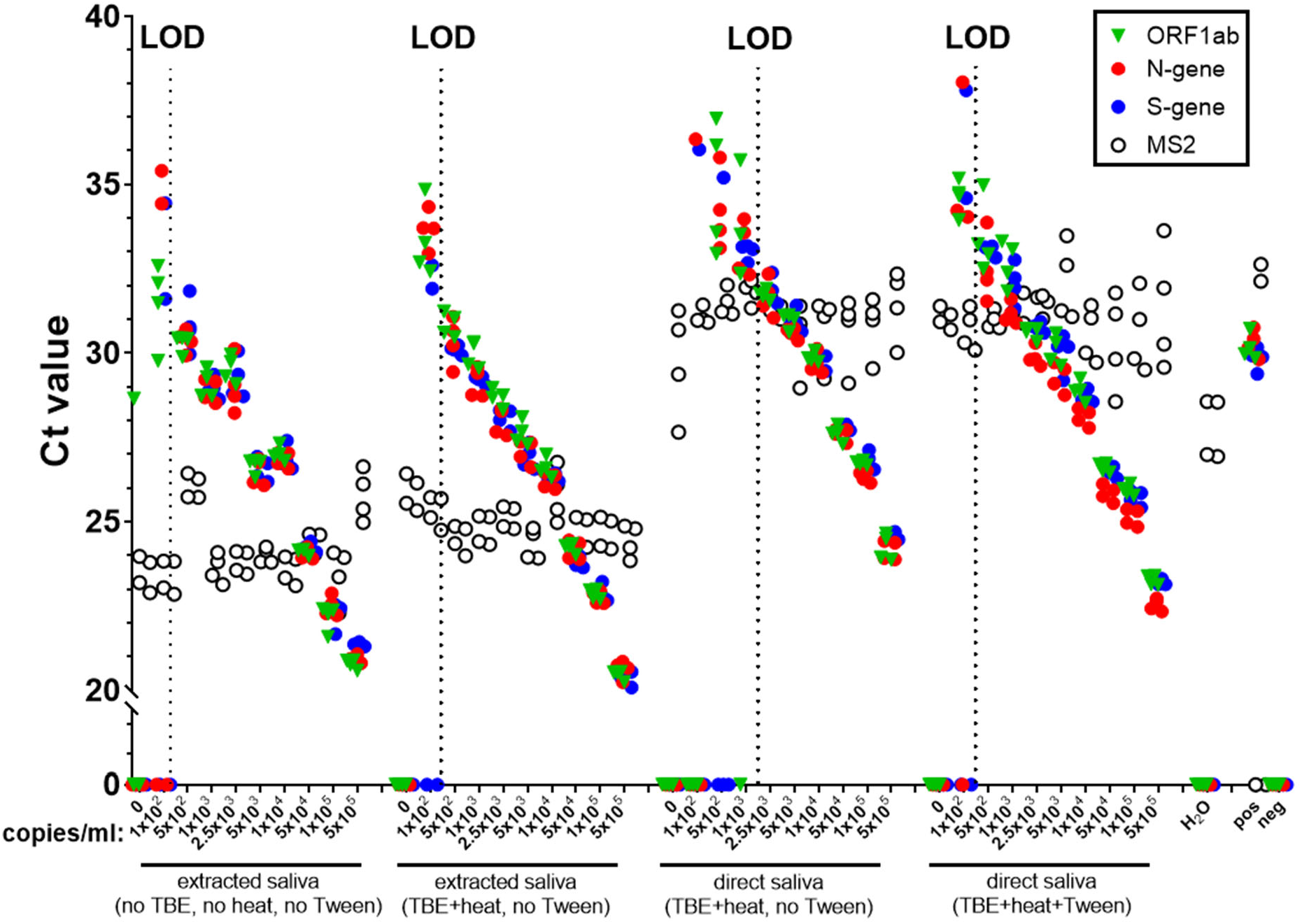
Limit of Detection (LOD) for assessment of SARS-CoV-2 from saliva, comparing a process utilizing RNA isolation/purification to one that bypasses RNA isolation/purification. γ-irradiated SARS-CoV-2 was spiked into fresh human saliva (SARS-CoV-2 negative), with or without TBE buffer(1X) at 1.0×10^2^, 5.0×10^2^, 1.0×10^3^, 2.5×10^3^, 5.0×10^3^, 1.0×10^4^, 5.0×10^4^, 1.0×10^5^, and 5.0×10^5^ viral copies/mL. Samples were incubated at 95°C for 30 min, then combined with or without Tween 20 (0.5%). All saliva samples were spiked with purified MS2 bacteriophage (1:40 MS2:sample) as an internal control. Virus-spiked saliva samples were either processed for RNA extraction followed by RT-qPCR (purified RNA), or directly analyzed by RT-qPCR (direct saliva). All samples, including a positive control (pos; SARS-CoV-2 positive control, 5.0×10^3^ copies/mL, no MS2) and a negative control (neg; water, no MS2) were analyzed by RT-qPCR, in triplicate, for SARS-CoV-2 ORF1ab (green triangle), N-gene (red square), and S-gene (blue circle), and MS2 (open circle). Undetermined Ct values are plotted at 0. The limit of detection (LOD) is indicated by the dotted vertical line.

As the TaqPath/MasterMix RT-qPCR reagents from ThermoFisher provide the necessary specificity for SARS-CoV-2 detection in a simplified workflow, this system was utilized for all the experiments described above. However, we have also assessed the Centers for Disease Control and Prevention (CDC)-approved primers and probes for SARS-CoV-2 N1 and N2 genes, and the human RNase P (RP) gene control in this direct saliva-to-RT-qPCR protocol, and the results show that these primers give comparable LOD values, with 5000 viral copies/mL using heat-inactivated SARS-CoV-2, and 500 viral copies/mL using γ-irradiated SARS-CoV-2 (**Supporting Figure 5**). These findings further illustrate that our optimized protocol may be used with comparable detection across multiple analytical platforms. Altogether, these findings indicate that the optimized protocol (heat treatment of saliva samples at 95°C for 30 min / addition of TBE buffer and Tween 20) yields a LOD that is comparable to reported clinical viral shedding concentrations in oral fluid, thus emphasizing the translatability of the protocol to detecting SARS-CoV-2 in patient samples.

#### Sample handling optimization

In preparation for clinical samples and real-world testing, we first evaluated the ability to detect spiked inactivated virus in samples that were stored at varying temperatures (ambient (25°C), 4°C, −20°C, and −80°C), for varying lengths of time (≤24 hrs). Most importantly, at room temperature and at 4°C samples processed after 1 hr showed little difference from those processed after 24 hr storage, suggesting considerable flexibility in processing time (**Supporting Figure 6**). Some increased variability between individual gene replicates and loss of signal was observed with prolonged storage and freeze/thaw cycles (**Supporting Figure 6**).

Next, evaluation was made of the effect of sample volume in the saliva collection vessels (50 mL conical tubes) on viral detection, after heating at 95°C for 30 min in a hot water bath, due to concerns of evaporation of smaller samples and incomplete heating of larger samples. No appreciable difference was observed across the anticipated range of clinical saliva sample volumes (0.5-5 mL), indicating that sample volume does not impact virus detection (**Supporting Figure 7**). Furthermore, if samples are transferred to smaller vessels for more efficient long-term cold storage (1.5 mL microcentrifuge tubes), no appreciable differences in virus detection between different volumes is anticipated (**Supporting Figure 7**). Finally, as clinical saliva samples can sometimes contain particulates, we next evaluated whether removal of the particulates via centrifugation affected viral detection (**Supporting Figure 8**). Notably, if samples were centrifuged, with the resultant supernatant being used for direct RT-qPCR, the LOD was approximately 10-fold worse, with fewer individual gene replicates being detected at lower viral copy numbers (**Supporting Figure 8**). Therefore, we recommend avoiding centrifugation of samples if possible. Altogether, these findings suggest that (1) saliva samples are stable under varying storage conditions, (2) the volume of sample heated with collection vessels does not affect viral detection, and (3) centrifugation of samples should be avoided for direct saliva-to-RT-qPCR testing of SARS-CoV-2.

#### LOD reproducibility

In order to evaluate the robustness of the optimized direct saliva-to-RT-qPCR approach, the LOD of 1000 SARS-CoV-2 viral copies/mL was measured in 30 independent replicate samples (**Figure 5**). γ-irradiated SARS-CoV-2 was spiked into fresh saliva from two healthy donors, and two commercially available saliva sources. Across all replicates, these samples with 1000 viral copies/mL were consistently detected (all three viral genes), further testifying to the ability of direct saliva-to-RT-qPCR to detect SARS-CoV-2. In order to validate the specificity of our detection system to SARS-CoV-2, saliva was spiked with or without SARS-CoV-2 (γ-irradiated virus, synthetic N-transcript), two other human coronaviruses (OC43, 229E), SARS and MERS synthetic RNA, and human RNA (extracted from HEK 293 cells). Among these samples, SARS-CoV-2 genes were only detected in the positive control, and SARS-CoV-2 samples, further supporting the specificity of the detection platform for SARS-CoV-2 (**Supporting Figure 9**).

**Figure 5.**
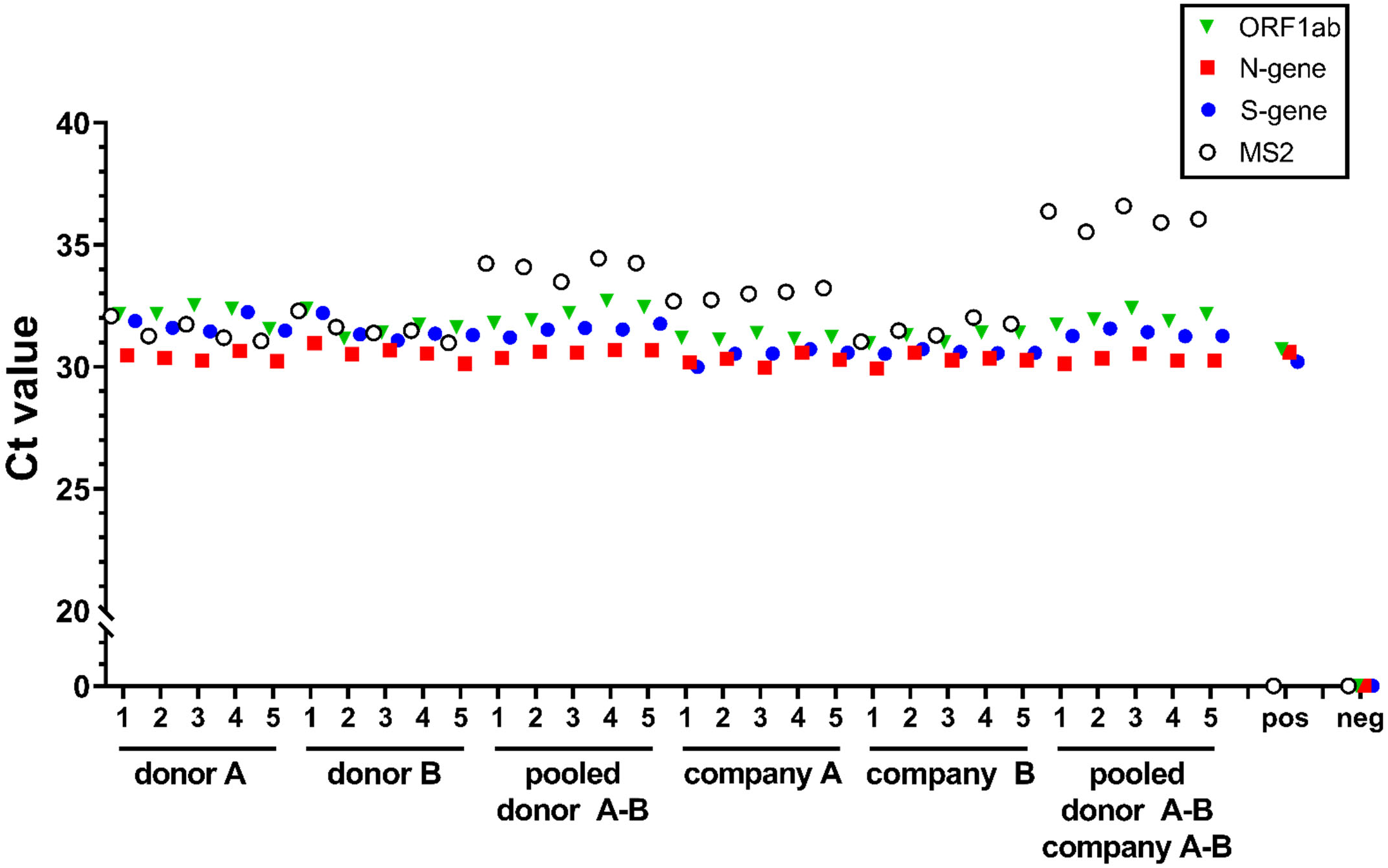
Limit of Detection (LOD) reproducibility. γ-irradiated SARS-CoV-2 was spiked into human saliva (SARS-CoV-2 negative), sourced fresh from two healthy donors, and purchased from two companies, in 1X TBE buffer at 1.0×10^3^ viral copies/mL. Samples were incubated at 95°C for 30 min, then Tween 20 was added to a final concentration of 0.5%. All saliva samples were spiked with purified MS2 bacteriophage (1:40 MS2:sample) as an internal control. Virus-spiked saliva samples were directly analyzed by RT-qPCR (direct saliva). All samples, including a positive control (pos; SARS-CoV-2 positive control, 5.0×10^3^ copies/mL, no MS2) and a negative control (neg; water, no MS2) were analyzed by RT-qPCR, in replicates of 5, for SARS-CoV-2 ORF1ab (green triangle), N-gene (red square), and S-gene (blue circle), and MS2 (open circle). Undetermined Ct values are plotted at 0.

### Clinical validation of direct saliva-to-RT-qPCR for diagnosis of SARS-CoV-2

Our findings support an optimized SARS-CoV-2 diagnostic approach that increases accessibility to testing by using saliva (rather than NP swabs) and eliminates the need for RNA extraction (thus saving time and resources). We next sought to assess our protocol with clinical samples. Although the changes in viral load in the NP cavity and in saliva over time are unknown, there is reason to believe they are different,^40,41^ so exact concordance between the two samples might not be expected; detection in saliva can provide complementary information to that in the NP cavity.

To evaluate the ability of the direct saliva-to-RT-qPCR approach to detect SARS-CoV-2 in clinical patient specimens, saliva was collected contemporaneously with NP swabs from 100 individuals using the following protocol: After saliva collection, TE was added at a 1:1 ratio, and samples were frozen for over a week before processing. For the evaluation, samples were thawed, 10X TBE buffer was added to a final concentration of 1X, heated at 95°C for 30 min, cooled to room temperature, and Tween 20 was added to a final concentration of 0.5%, followed by direct RT-qPCR. Given biological complexity in clinical samples, variabilities in signal detection based on viral load and gene target length (ORF1ab > S > N) may occur; therefore, a given result was interpreted as positive if one or more gene targets were detected, and negative if no gene targets were detected. Furthermore, a result was considered valid if all gene targets were detected in the SARS-CoV-2 positive control and no gene targets were detected in the negative control. A notable power in the context of a multiplex system is the ability to evaluate three independent viral genes in a single reaction, rather than relying upon multiple probes across different reactions for a single viral gene (as is used in other systems). One of the benefits of saliva-based testing is the possibility of frequent and easy retesting of samples and of individuals, and as such duplicate testing (testing of the same saliva sample two different times) was utilized for this study.

Of the 100 samples analyzed, 9 were positive for SARS-CoV-2 as assessed by NP swab, and upon duplicate testing the direct saliva-to-RT-qPCR process identified the same 9 samples as positive, with 8 of 9 saliva samples positive in both of the replicates. All 91 samples identified as negative by NP swab were also negative via saliva testing, although in one of these samples one of the duplicate runs was positive, but was negative upon re-tests (**Figure 6, Table 1**). Even though these samples were not run under the fully optimized protocol, this initial testing of clinical samples using direct saliva-to-RT-qPCR showed excellent performance. When testing samples a single time, it was 88.9% sensitive and 98.9% specific for SARS-CoV-2, with an 11.1% false negative and 1.1% false positive rate, and 88.9% positive and 98.9% negative predictive value. Using duplicate testing of samples, sensitivity and specificity, and positive and negative predictive values, all increased to 100%, and the false negative and positive rates decreased to 0%.

**Table 1.**
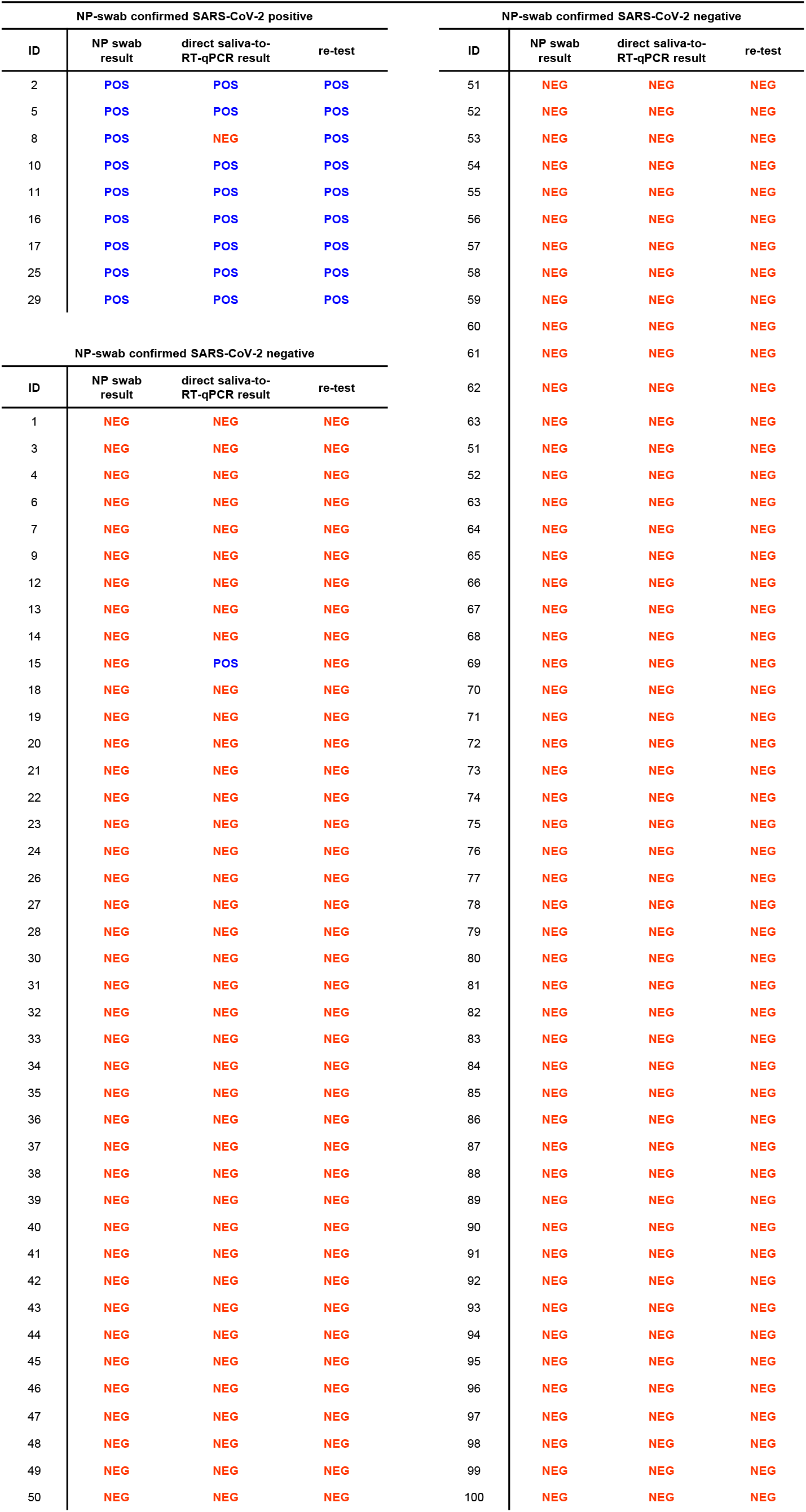
Clinical evaluation of direct saliva-to-RT-qPCR.

**Figure 6.**
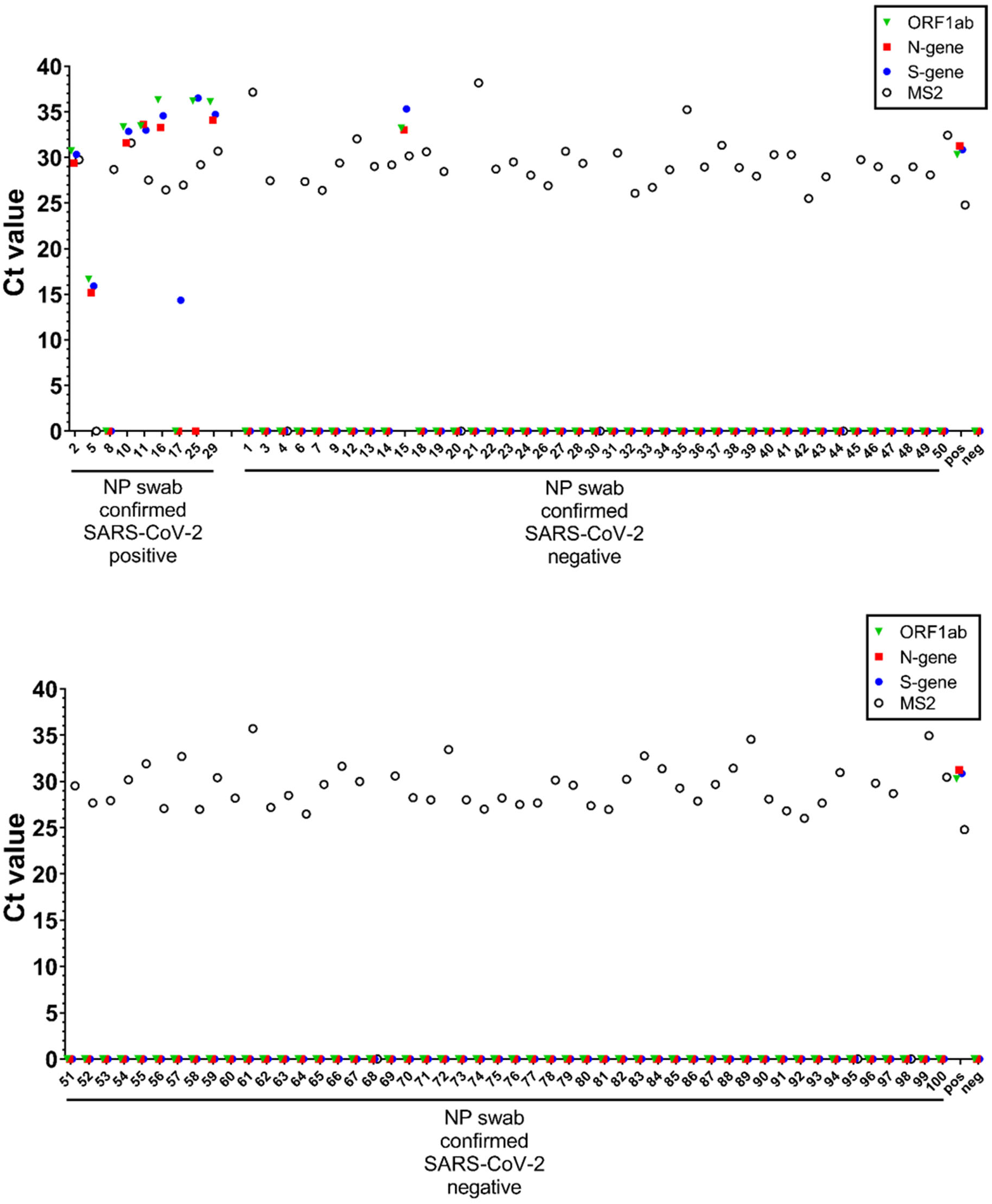
Assessment of clinical samples. Saliva samples from 9 SARS-CoV-2 positive and 91 SARS-CoV-2 negative patients (as judged by NP swabs in VTM with RNA extraction) had TE buffer added to them (at a 1:1 ratio) and were frozen for over a week. Upon thawing, 10X TBE buffer was added to the samples at a final concentration of 1X, heated at 95°C for 30 min, cooled to room temp, and Tween 20 was added to a final concentration of 0.5%. All saliva samples were spiked with purified MS2 bacteriophage (1:40 MS2:sample) as an internal control. Saliva samples were directly analyzed by RT-qPCR. All samples, including a positive control (pos; SARS-CoV-2 positive control, 5.0×10^3^ copies/mL) and a negative control (neg; water) were analyzed by RT-qPCR, in singlet, for SARS-CoV-2 ORF1ab (green triangle), N-gene (red square), and S-gene (blue circle), and MS2 (open circle). This figure shows one of the two replicates. Undetermined Ct values are plotted at 0.

## Discussion

### Comparison of NP swab and saliva-based testing

When seeking to develop a SARS-CoV-2 molecular diagnostic protocol suitable for testing >10,000 individuals a day, the ease with which saliva can be collected, and the known presence of the virus in saliva makes it highly desirable as the sample medium. As a diagnostic tool, such testing has the additional advantage of making assessments directly from an oral fluid that may be a culprit in transmission of SARS-CoV-2.^10^ Unfortunately, only a handful of studies have examined the viral load dynamics over time for saliva and NP swab samples.^40,41^ While these studies support the notion that SARS-CoV-2 tends to be at its highest level in saliva during the first week of infection, more information is needed on this important topic. In contrast, studies have shown that while live virus can no longer be cultured from patients 10 days after symptom onset,^42^ NP swabs continue to be positive after a patient is in the convalescent phase and no longer infectious.^13^ As such, it is quite possible that differences observed in studies comparing SARS-CoV-2 levels in saliva and NP swabs are real, and not an artifact of different testing sensitivities; while in general concordance between the NP swab and saliva testing has been high in other studies (87%,^40^ 92%,^14^ 100%^43^), results will likely depend on what point during infection a patient is sampled.

### Direct saliva-to-RT-qPCR process, key advances and remaining limitations

The direct saliva-to-RT-qPCR method described herein, bypassing NP swabs, VTM, and RNA isolation/purification, was enabled by a handful of key discoveries. First, the time and duration of heating the saliva sample is critical. Standard protocols for heat inactivation of SARS-CoV-2 call for heating at ~60°C for 30 minutes;^30,31^ while these conditions inactivate the virus, they do not allow for successful SARS-CoV-2 detection via direct RT-qPCR, likely because of the persistence of as-yet-unidentified factors in saliva that are inhibitory to RT-qPCR. Heating at 95°C for 30 minutes likely inactivates these inhibitory components and allows for excellent SARS-CoV-2 detection in this direct process that bypasses RNA isolation/purification. Second, while TE buffer performs well, consistent with another report successfully using TE to extract dry NP swabs,^17^ TBE buffer provides more reliability and consistency in our direct saliva-to-RT-qPCR detection of SARS-CoV-2. Finally, the addition of the non-ionic detergent Tween-20 also helped improve detection of SARS-CoV-2, possibly by facilitating the opening of the viral capsid to allow the release of RNA to provide sufficient template for RT-qPCR detection.

Our preliminary assessment of clinical samples is very promising, especially given that these samples were not collected and processed under the optimized protocol (they were collected before our discovery of the benefits of TBE buffer and Tween 20); with these samples TE buffer was added to the sample, and they were frozen for over a week before processing. However, even under this non-optimized workflow we were able to identify all 9 NP swab positives with duplicate runs of the samples. Next steps are to perform similar head-to-head comparisons between the NP swab-based method and our optimized workflow with additional clinical samples.

### Supply chain, costs, and next-generation technology

A major benefit of the simple workflow (see Graphical Abstract) detailed herein is its ability to be adopted by any diagnostic laboratory currently using RT-qPCR in SARS-CoV-2 testing. In addition to the time savings and major logistical benefits of using saliva and bypassing RNA isolation/purification, our analysis of the costs of all reagents/disposables for this process amounts to ~$10 per test, the bulk of which are the TaqPath/MasterMix. This cost could drop further if samples are pooled before RT-qPCR. Pooling considerations will necessarily be informed by data on the expected positive rate in the population to be tested, and also the relationship between viral load and infectivity;^44–46^ while one recent study showed that live SARS-CoV-2 could not be cultured from samples containing less than 1,000,000 viral copies per mL,^42^ more information is needed. And, while there is no indication that TaqPath/MasterMix will be limited by the supply chain, we show that this process and workflow is also compatible with other primer sets, such as the N1 and N2 primers and probes from the CDC. In the future, development of analogous saliva-based processes that bypass RNA isolation/purification can be envisioned for alternative back-end detection technologies, such as the LAMP method,^2,47^ which if successful would result in an even shorter overall time from sample collection to results.

In summary, described herein is a sensitive diagnostic method for SARS-CoV-2 that is operationally simple, bypasses supply chain bottlenecks, evaluates a clinically relevant infectious fluid, is appropriate for large scale repeat testing, is cost effective, and can be readily adopted by other laboratories. Large scale SARS-CoV-2 testing will be a powerful weapon in preventing spread of this virus and helping to control the COVID-19 pandemic.

## Acknowledgement

This work was supported by the University of Illinois, Urbana-Champaign. We would like to thank Dr. Rashid Bashir and Dr. Enrique Valera for their assistance in coordinating and obtaining IRB approval for acquisition of clinical samples, as well as Dr. Mark Johnson, Reubin McGuffin, MaryEllen Sherwood, and Carly Skadden for their assistance in collection and distribution of saliva and discarded VTM samples from Carle Foundation Hospital. We are grateful to Dr. Lois Hoyer for use of QuantStudio3 RT-qPCR instruments.

## Materials and Methods

### Acquisition and processing of clinical samples

All clinical samples from study participants were collected in accordance with University of Illinois at Urbana-Champaign (UIUC) IBC-approved protocol number 4604 and IRB-approved protocol number 20CRU3150. Saliva in 1:1 1X TE buffer and discarded VTM samples collected from 100 adults at the Carle Foundation Hospital Drive-thru COVID-19 testing center were collected and frozen at −80°C for over a week. Upon thawing, 10X TBE buffer was added to the samples to a final concentration of 1X, heated at 95°C for 30 min, cooled to room temperature, and Tween 20 was added to a final concentration of 0.5%. The optimized direct saliva-to-RT-qPCR approach was compared to detection of SARS-CoV-2 from nasopharyngeal (NP) swab in VTM performed at the Carle Foundation Hospital. In all studies conducted, researchers were blinded to the results obtained from clinical RT-qPCR tests performed on NP swabs at the Carle Foundation Hospital.

### Collection and processing of fresh saliva from healthy donors

Fresh saliva was collected from healthy individuals in 50 mL conical tubes (BD Falcon) in accordance with University of Illinois at Urbana-Champaign IBC-approved protocol numbers 4604 and 4589. In some experiments, pooled saliva from healthy donors was purchased from Lee BioSolutions, Inc. (CN 991-05-P) and Innovative Research (CN IRHUSL50ML). Saliva was diluted at a 1:1 ratio with either TBE buffer (100mM Tris-HCl pH8.0, 90mM boric acid, and 1mM EDTA) or TE buffer (10mM Tris-HCl pH8.0 and 1mM EDTA) buffer. In some experiments, Phosphate Buffered Saline (PBS), DNA/RNA Shield (Zymo Research), and SDNA-1000 (Spectrum Solutions), were also tested at final working concentrations of 2X, 1.5X, 1X, and 0.5X. Known amounts of the SARS-CoV-2 inactivated virus (BEI) were spiked into saliva samples. Samples were incubated in a hot water bath at 95°C for 30 min. All saliva samples were spiked with purified MS2 bacteriophage (1:40 MS2:sample) as an internal control. In some experiments, RNA extraction was performed on 200 μL saliva (+/- virus) using MagMax Viral/Pathogen II Nucleic Acid Isolation Kit (Applied Biosciences CN A48383) following the manufacturer’s protocol. Extracted RNA was eluted from magnetic beads in 50μl UltraPure DNase/RNase-free distilled water (Ambion CN 10977023). RNA concentration of eluted RNA was measured using Qubit RNA Broad Range (BR) assay kit (Fisher Scientific).

### SARS-CoV-2 inactivated virus and human coronaviruses

In most experiments, fresh pooled saliva diluted 1:1 in TBE buffer (1x final concentration) were spiked with either gamma-irradiated (BEI cat# NR-52287, Lot no. 70033322) or heat-inactivated (BEI cat# NR-52286, Lot no. 70034991) SARS-CoV-2 virions. The reported genome copy number pre-inactivation for γ-irradiated and heat-inactivated SARS-CoV-2 are 1.7×10^9^ and 3.75×10^8^ genome equivalents/mL, respectively, for the specified lot numbers. The following reagent was deposited by the Centers for Disease Control and Prevention and obtained through BEI Resources, NIAID, NIH: SARS-Related Coronavirus 2, Isolate USA-WA1/2020, Gamma-irradiated, NR-52287, and heat-inactivated, NR-52286. Seasonal human coronaviruses (OC43 and 229E strains) were obtained from the World Reference Center for Emerging Viruses and Arboviruses at UTMB.

Genomic RNA for SARS-Related Coronavirus 2 (Isolate USA-WA1/2020), NR-52285, was obtained from BEI Resources. In addition, the 2019-nCoV_N_Positive Control (CN 10006625), SARS-CoV Control (CN 10006624), and MERS-CoV Control (CN 10006623) synthetic RNA transcripts were purchased from Integrated DNA Technologies.

All virus stocks and RNA transcripts were aliquoted in small volumes and stored at −70°C. Stocks were serially diluted to the correct concentration in RNase-free water on the day of experimentation.

### RT-qPCR assay

We performed a multiplex RT-qPCR assay using the TaqPath RT-PCR COVID-19 kit (Thermo Fisher CN A47814) together with the TaqPath 1-step master mix – No ROX (Thermo Fisher CN A28523). To reduce cost, RT-qPCR reactions were prepared at half the suggested reaction mix volume (7.5 μL instead of 15 μL). 10 μL of either saliva in TBE buffer or extracted RNA were used as templates for the RT-qPCR reaction. All saliva samples used for pre-clinical studies were spiked with purified MS2 bacteriophage (1:40 MS2:sample) as an internal control prior to analysis by RT-qPCR. For clinical samples, MS2 was added to the preparation of the reaction mix (1μL MS2 per reaction). COVID-19 positive control RNA at 25 genomic copies/μL was used. Negative control is UltraPure DNase/RNase-free distilled water (Ambion CN 10977023). All RT-qPCR reactions were performed in 0.2 mL 96-well reaction plates in a QuantStudio 3 system (Applied Biosciences). The limit of detection (LOD) of the assay was performed by serial dilution of γ-irradiated SARS-CoV-2 (0-5.0×10^5^ viral copies/mL) used to spike pooled fresh saliva samples. The RT-qPCR was run using the standard mode, consisting of a hold stage at 25°C for 2 min, 53°C for 10 min, and 95°C for 2 min, followed by 40 cycles of a PCR stage at 95°C for 3 sec then 60°C for 30 sec; with a 1.6°C/sec ramp up and ramp down rate.

In some experiments, the CDC-approved assay was used to validate our data using the TaqPath 1-step mix (Thermo Fisher CN A15300). Primers and probes targeting the N1, N2, and RP genes were purchased from Integrated DNA Technologies as listed: nCOV_N1 Forward Primer Aliquot (CN 10006830), nCOV_N1 Reverse Primer Aliquot (CN 10006831), nCOV_N1 Probe Aliquot (CN 10006832), nCOV_N2 Forward Primer Aliquot (CN 10006833), nCOV_N2 Reverse Primer Aliquot (CN 10006834), nCOV_N2 Probe Aliquot (CN 10006835), RNase P Forward Primer Aliquot (CN 10006836), RNase P Reverse Primer Aliquot (CN 10006837), RNase P Probe Aliquot (CN 10006838). The 2019-nCoV_N_Positive Control (IDT CN 10006625) was used as positive control at 50 copies/μL dilution.

### Detergent optimization

γ-irradiated SARS-CoV-2 (1.0×10^4^ viral copies/mL) was spiked into fresh human saliva (SARS-CoV-2 negative) and combined 1:1 with Tris-Borate-EDTA buffer (TBE) at a final working concentration of 1X.

Samples were treated with varying concentrations of detergents (Triton X-100 (Fisher Scientific), Tween 20 (Fisher Scientific), NP-40 (Fisher Scientific)) before or after heating at 95°C for 30 min. All saliva samples were spiked with purified MS2 bacteriophage (1:40 MS2:sample) as an internal control prior to analysis by RT-qPCR (Fisher TaqPath COVID-19 Combo kit, QuantStudio 3).

### Sample volume heat treatment optimization

γ-irradiated SARS-CoV-2 (1.0×10^4^ viral copies/mL; BEI) was spiked into fresh human saliva (SARS-CoV-2 negative) and combined 1:1 with Tris-Borate-EDTA buffer (TBE) at a final working concentration of 1X. The sample was distributed into either 50 mL conical (BD Falcon) or 1.5 mL microfuge tubes (Ambion), at either 10% (5 mL in 50 mL conical, 150 μL in 1.5 ml microfuge), 5% (2.5 ml in 50 mL conical, 75 μL in 1.5 mL microfuge), or 1% (0.5 mL in 50 mL conical, 15 μL in 1.5 mL microfuge) the vessel storage capacity. Samples were incubated in a hot water bath at 95°C for 30 min. All saliva samples were spiked with purified MS2 bacteriophage (1:40 MS2:sample) as an internal control prior to analysis by RT-qPCR (Fisher TaqPath COVID-19 Combo kit, QuantStudio 3).

### Sample buffer additive optimization

γ-irradiated SARS-CoV-2 (1.0×10^4^ viral copies/mL) was spiked into fresh human saliva (SARS-CoV-2 negative) and combined 1:1 with Tris-Borate-EDTA buffer (TBE) at a final working concentration of 1X in 50 mL conical tubes (BD Falcon). Samples (1.0 mL in 50 mL conical tubes) were incubated in a hot water bath at 95°C for 30 min. Following heat treatment, virus-spiked saliva was aliquoted in 1.5 mL tubes and combined with various RNA stabilizing agents to a final volume of 40 μL. Additives include RNaseI (1 U/μL), carrier RNA (0.05 μg/mL), glycogen (1 μg/μL), TCEP/EDTA (1X), Proteinase K (5 μg/μL), RNase-free BSA (1.25 mg/ml), RNAlater (1:1 ratio in place of TBE), or PBS-DTT (6.5 mM DTT in PBS, diluted 1:1 in place of TBE). All saliva samples were spiked with purified MS2 bacteriophage (1:40 MS2:sample) as an internal control prior to analysis by RT-qPCR (Fisher TaqPath COVID-19 Combo kit, QuantStudio 3).

### Saliva stability optimization

Pre-aliquoted γ-irradiated SARS-CoV-2 (1.0×10^4^ viral copies/mL) was spiked into pre-aliquoted fresh human saliva (SARS-CoV-2 negative) and combined with Tris-Borate-EDTA buffer (TBE), at a final working concentration of 1X. Samples (0.5 mL in 50 mL conical tubes) were stored at 25°C (ambient temperature), 4°C, −20°C, or −80°C for 1, 2, 4, 8, 12, and 24 hours. All saliva samples were spiked with purified MS2 bacteriophage (1:40 MS2:sample) as an internal control prior to analysis by RT-qPCR (Fisher TaqPath COVID-19 Combo kit, QuantStudio 3).

### Data analysis

Following completion of RT-qPCR, data were processed using QuantStudio Design and Analysis Software (version 1.2). Cycle threshold (Ct) values were plotted as single replicate values on a scatter plot, using GraphPad Prism 8 (version 8.4.2). Sensitivity, specificity, false positive, false negative, positive predictive values, and negative predictive values were calculated using the current standard for SARS-CoV-2 detection (NP swabs in VTM with RNA extraction) as confirmation of true disease positive and disease negative status.

## Supporting Figures

**Supporting Figure 1.**
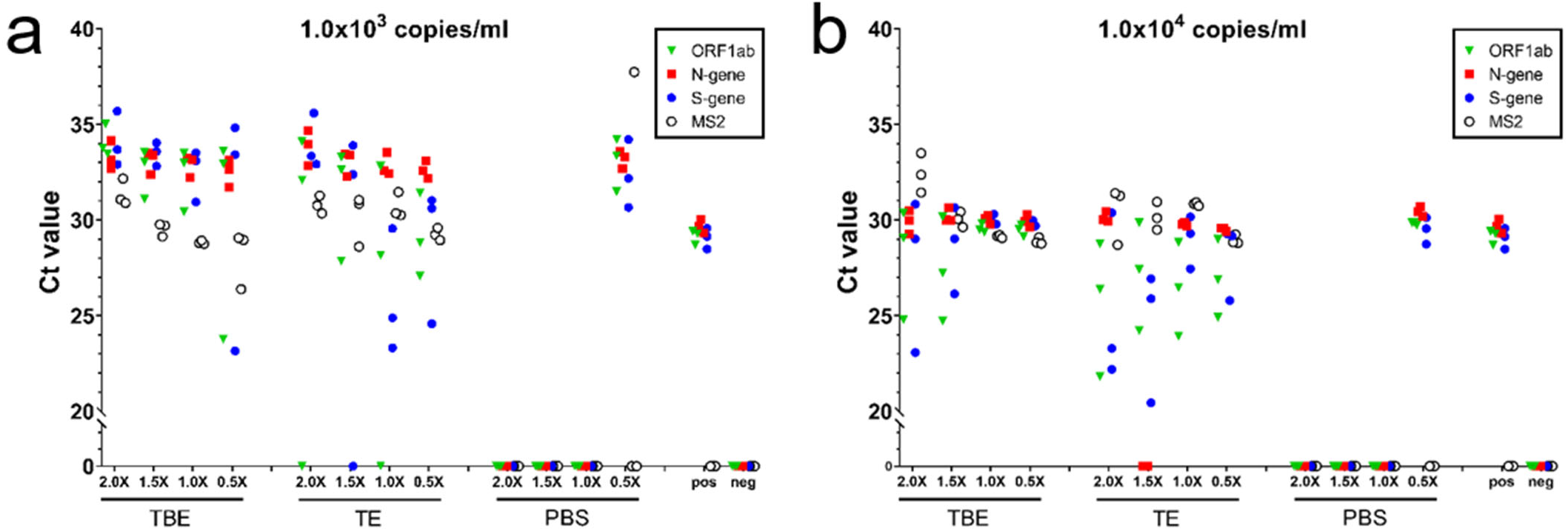
Saliva collection buffer titration. γ-irradiated SARS-CoV-2 (1.0×10^3^ (a) or 1.0×10^4^ (b) viral copies/mL) was spiked into fresh human saliva (SARS-CoV-2 negative) and combined with Tris-Borate-EDTA buffer (TBE), Tris-EDTA buffer (TE), or Phosphate Buffered Saline (PBS), at a final working concentration of 2X, 1.5X, 1X, or 0.5X. Samples (0.5 mL in 50 mL conical tubes) were incubated in a hot water bath at 95°C for 30 min. All saliva samples were spiked with purified MS2 bacteriophage (1:40 MS2:sample) as an internal control. Virus-spiked saliva samples, a positive control (pos; SARS-CoV-2 positive control, 5.0×10^3^ copies/mL, no MS2) and a negative control (neg; water, no MS2) were directly analyzed by RT-qPCR, in triplicate, for SARS-CoV-2 ORF1ab (green triangle), N-gene (red square), and S-gene (blue circle), and MS2 (open circle). Undetermined Ct values are plotted at 0.

**Supporting Figure 2.**
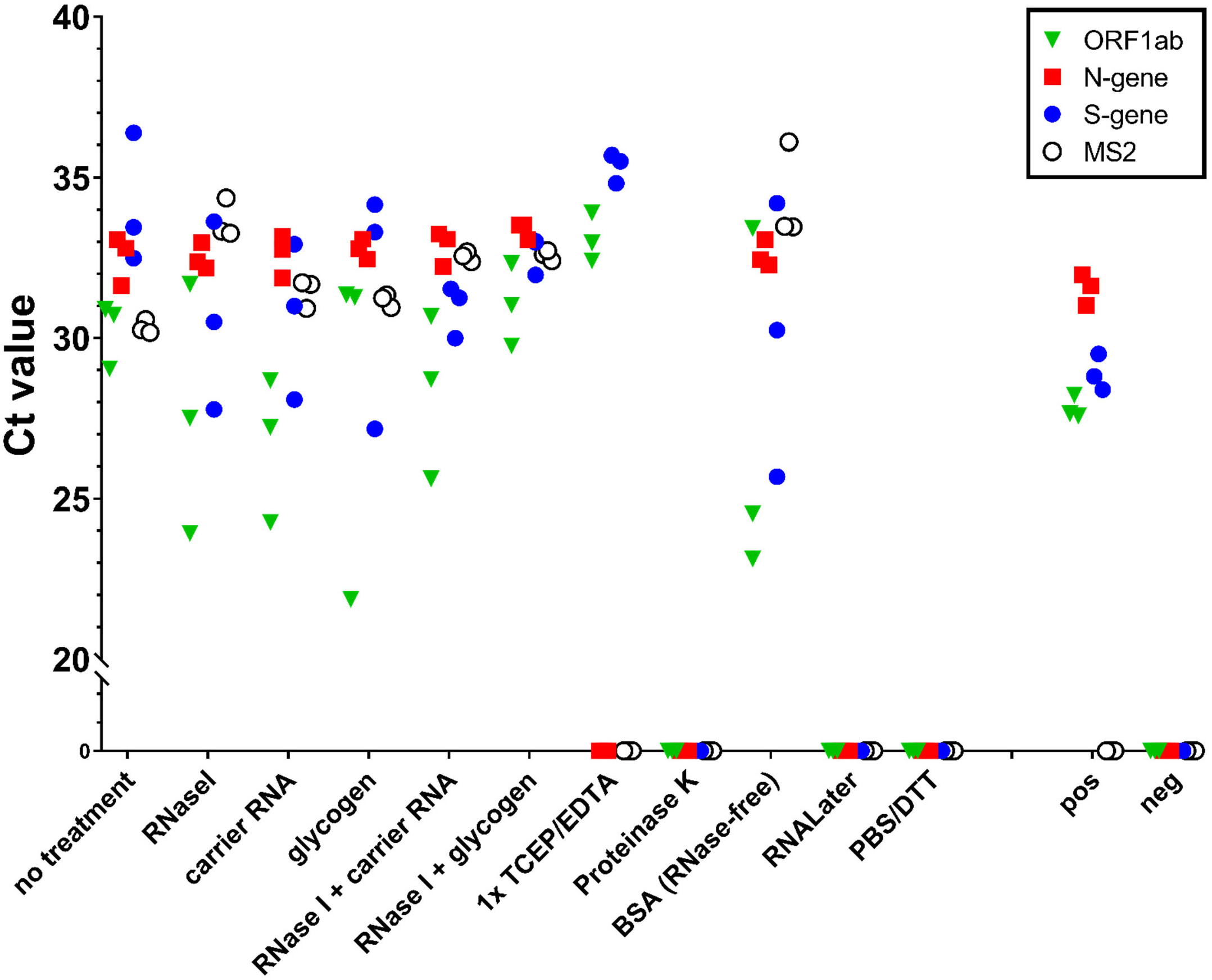
RNA stabilizing additive optimization. γ-irradiated SARS-CoV-2 (1.0×10^4^ viral copies/mL) was spiked into fresh human saliva (SARS-CoV-2 negative) and combined with TBE buffer, at a final working concentration of 1X. Samples (0.5 mL in 50 mL conical tubes) were incubated in a hot water bath at 95°C for 30 min. Following heat treatment, virus-spiked saliva was combined with various RNA stabilizing agents, including RNaseI (1 U/μL), carrier RNA (0.05 μg/mL), glycogen (1 μg/μL), TCEP/EDTA (1X), Proteinase K (5 μg/μL), RNase-free BSA (1.25 mg/mL), RNAlater (1:1 ratio in place of TBE), or PBS/DTT (6.5 mM DTT in PBS, diluted 1:1 in place of TBE). All saliva samples were spiked with purified MS2 bacteriophage (1:40 MS2:sample) as an internal control. Virus-spiked saliva samples with or without additives, a positive control (pos; SARS-CoV-2 positive control, 5.0×10^3^ copies/mL, no MS2) and a negative control (neg; water, no MS2) were directly analyzed by RT-qPCR, in triplicate, for SARS-CoV-2 ORF1ab (green triangle), N-gene (red square), and S-gene (blue circle), and MS2 (open circle). Undetermined Ct values are plotted at 0.

**Supporting Figure 3.**
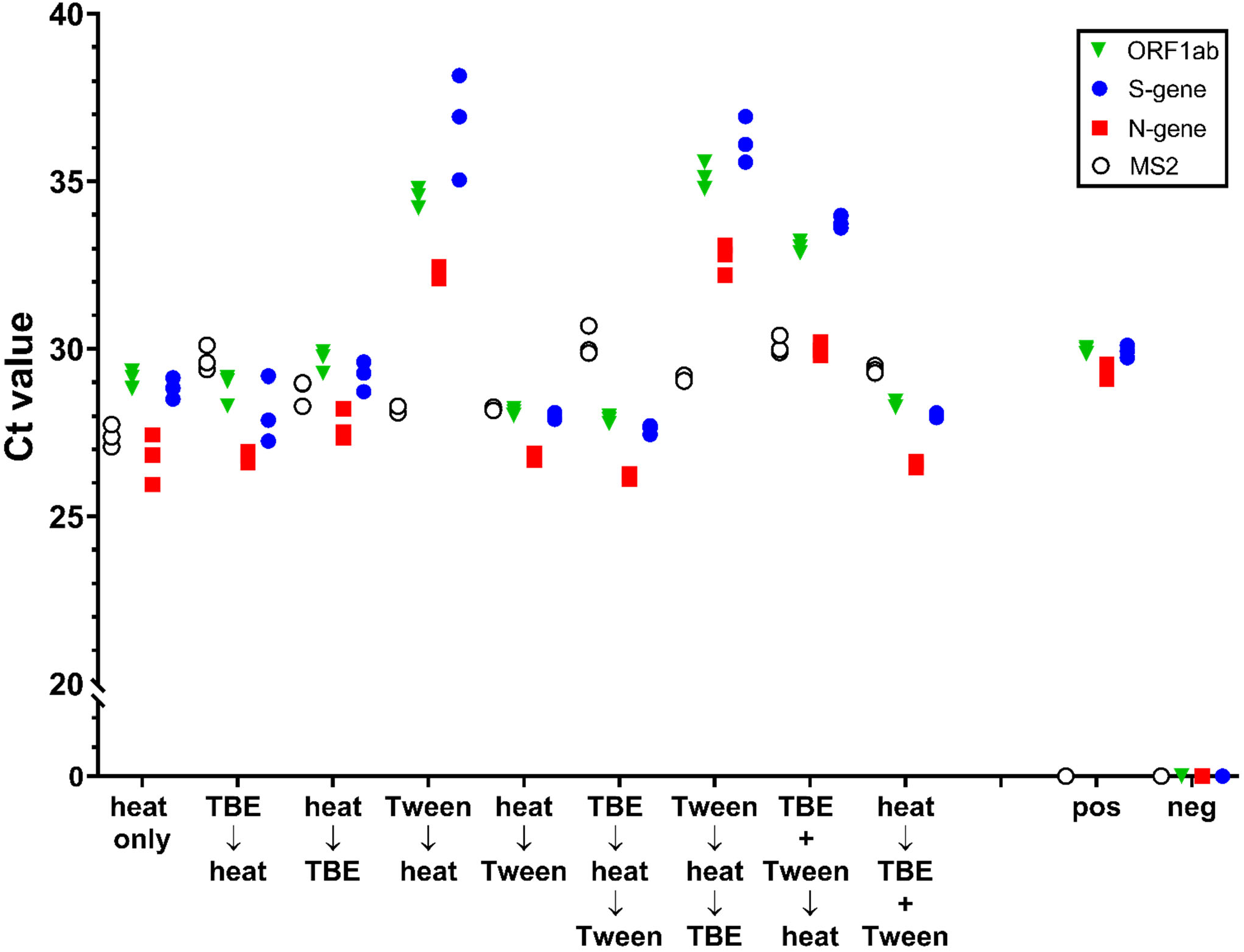
Workflow of TBE and Tween addition in relation to heat. γ-irradiated SARS-CoV-2 (1.0×10^5^ viral copies/mL) was spiked into fresh human saliva (SARS-CoV-2 negative) and combined with TBE buffer (1:10, final concentration 1X) and Tween 20 (1:20, final concentration 0.5%) alone or in combination, before or after heat treatment at 95°C for 30 min. All saliva samples were spiked with purified MS2 bacteriophage (1:40 MS2:sample) as an internal control. Virus-spiked saliva samples, a positive control (pos; SARS-CoV-2 positive control, 5.0×10^3^ copies/mL, no MS2) and a negative control (neg; water, no MS2) were directly analyzed by RT-qPCR, in triplicate, for SARS-CoV-2 ORF1ab (green triangle), N-gene (red square), and S-gene (blue circle), and MS2 (open circle). Undetermined Ct values are plotted at 0.

**Supporting Figure 4.**
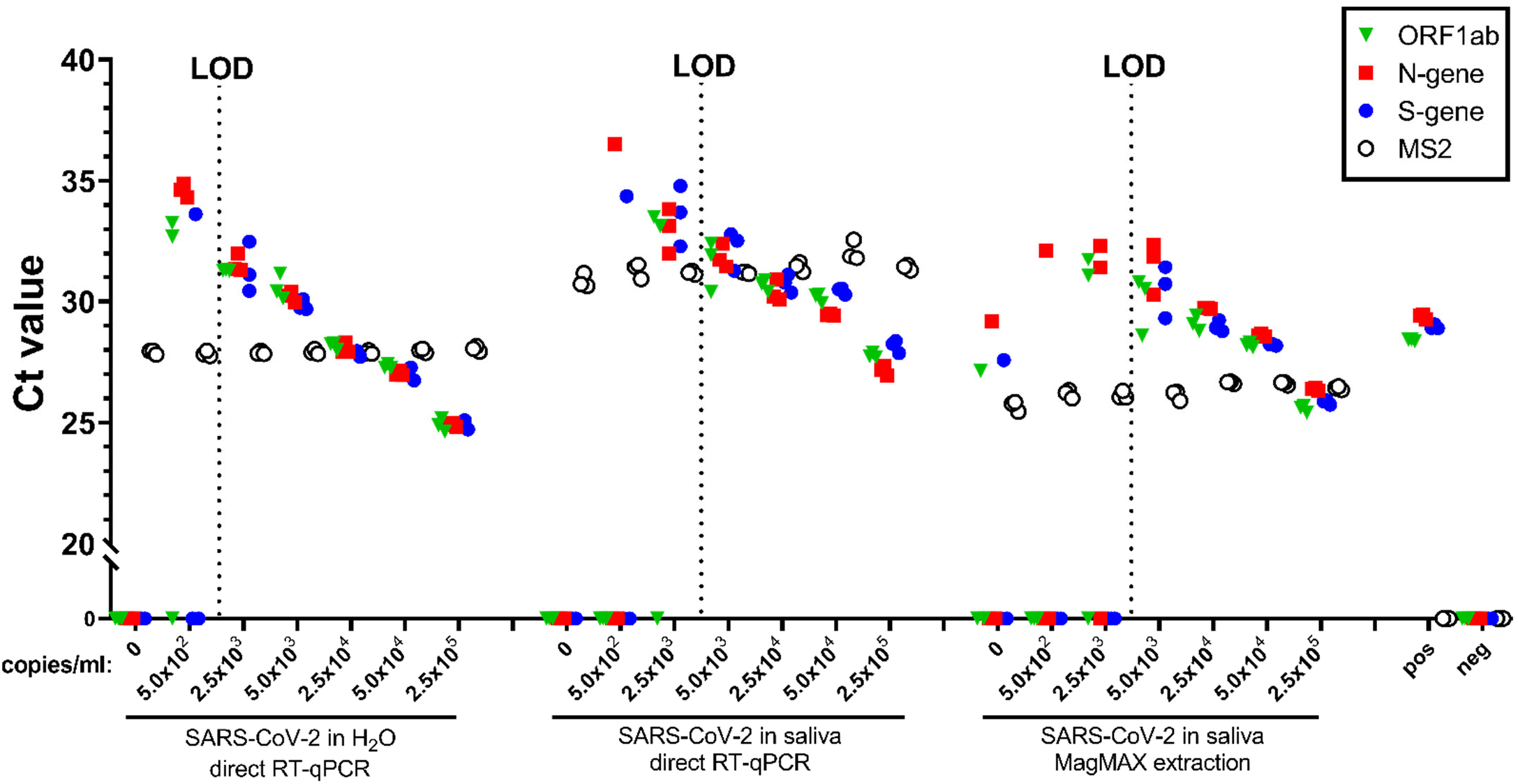
Limit of detection optimization. Heat-inactivated SARS-CoV-2 was spiked into fresh human saliva (SARS-CoV-2 negative) in 0.5X TE or water at 5.0×10^2^, 2.5×10^3^, 5.0×10^3^, 2.5×10^4^, 5.0×10^4^, and 2.5×10^5^ viral copies/mL. Samples were incubated at 95°C for 30 min. All samples were spiked with purified MS2 bacteriophage (1:40 MS2:sample) as an internal control. Virus-spiked samples were either processed for RNA extraction using a commercially available kit (MagMAX), or directly analyzed by RT-qPCR (direct saliva). All samples, including a positive control (pos; SARS-CoV-2 positive control, 5.0×10^3^ copies/mL, no MS2) and a negative control (neg; water, no MS2) were analyzed by RT-qPCR, in triplicate, for SARS-CoV-2 ORF1ab (green triangle), N-gene (red square), and S-gene (blue circle), and MS2 (open circle). Undetermined Ct values are plotted at 0. The limit of detection (LOD) is indicated by the vertical dotted line.

**Supporting Figure 5.**
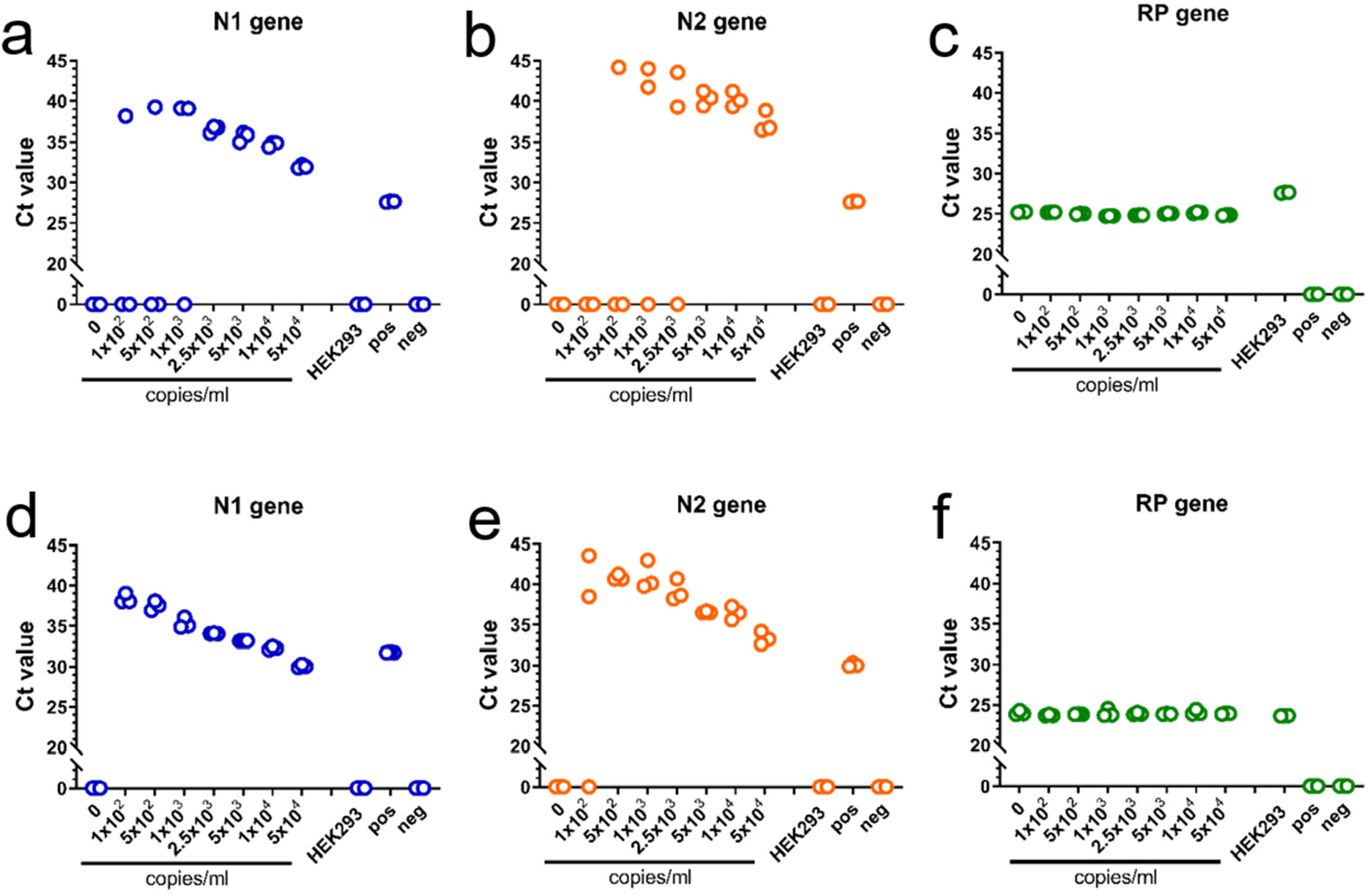
LOD of direct saliva-to-RT-qPCR SARS-CoV-2 detection using CDC-approved primers and probes. Heat-inactivated (a, b, c) and γ-irradiated (d, e, f) SARS-CoV-2 was spiked into fresh human saliva (SARS-CoV-2 negative) in 1X Tris-Borate-EDTA buffer (TBE) at 1.0×10^2^, 5.0×10^2^, 1.0×10^3^, 2.5×10^3^, 5.0×10^3^, 1.0×10^4^, and 5.0×10^4^ viral copies/mL. Samples were incubated at 95°C for 30 min. Virus-spiked saliva samples, a positive control (pos; SARS-CoV-2 positive control, 5.0×10^3^ copies/mL) and a negative control (neg; water) were directly analyzed by RT-qPCR, in triplicate, for SARS-CoV-2 N1 gene (a, d) and N2 gene (b, e), and the human RP gene (c, f). Undetermined Ct values are plotted at 0.

**Supporting Figure 6.**
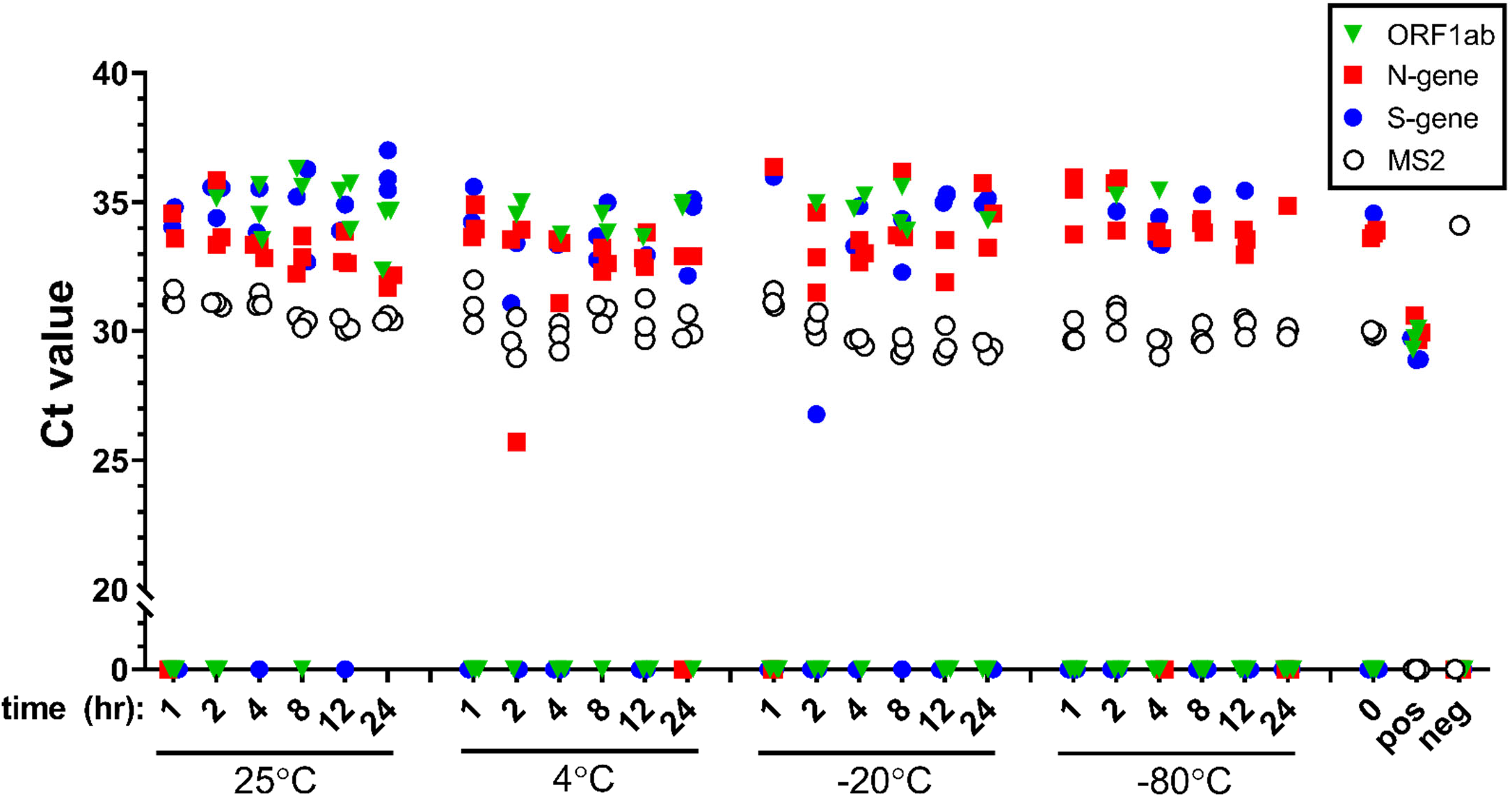
Stability of saliva samples. γ-irradiated SARS-CoV-2 (1.0×10^4^ viral copies/mL) was spiked into fresh human saliva (SARS-CoV-2 negative) and combined with TBE buffer 1:1 to a final working concentration of 1X. Samples (0.5 mL in 50 mL conical tubes) were stored at 25°C (ambient temperature), 4°C, −20°C, or −80°C for 1, 2, 4, 8, 12, and 24 hours. Following storage, samples were incubated in a hot water bath at 95°C for 30 min. All saliva samples were spiked with purified MS2 bacteriophage (1:40 MS2:sample) as an internal control. Virus-spiked saliva samples stored under different conditions, a freshly prepared virus-spiked saliva sample (0 hr), a positive control (pos; SARS-CoV-2 positive control, 5.0×10^3^ copies/mL, no MS2) and a negative control (neg; water, no MS2) were directly analyzed by RT-qPCR, in triplicate, for SARS-CoV-2 ORF1ab (green triangle), N-gene (red square), and S-gene (blue circle), and MS2 (open circle). Undetermined Ct values are plotted at 0.

**Supporting Figure 7.**
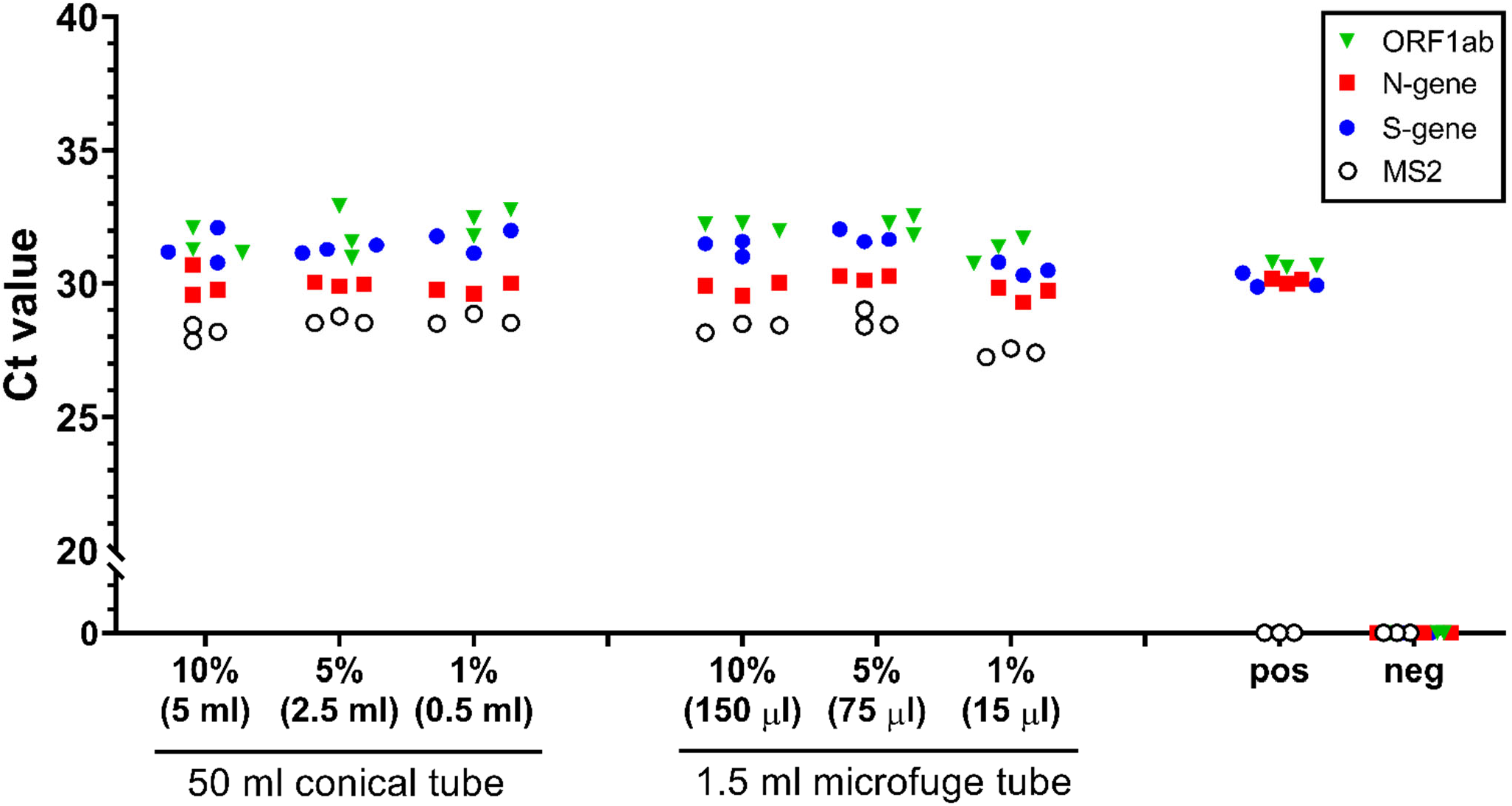
Effect of sample volume on SARS-CoV-2 detection. γ-irradiated SARS-CoV-2 (1.0×10^4^ viral copies/mL) was spiked into fresh human saliva (SARS-CoV-2 negative) and combined with TBE buffer 1:1 at a final working concentration of 1X. The sample was distributed into either 50 mL conical or 1.5 mL microfuge tubes, at either 10% (5 mL in 50 mL conical, 150 μL in 1.5 ml microfuge), 5% (2.5 mL in 50 ml conical, 75 μL in 1.5 ml microfuge), or 1% (0.5 mL in 50 mL conical, 15 μL in 1.5 mL microfuge) the vessel storage capacity. Samples were incubated in a hot water bath at 95°C for 30 min. All saliva samples were spiked with purified MS2 bacteriophage (1:40 MS2:sample) as an internal control. Virus-spiked saliva samples, a positive control (pos; SARS-CoV-2 positive control, 5.0×10^3^ copies/mL, no MS2) and a negative control (neg; water, no MS2) were directly analyzed by RT-qPCR, in triplicate, for SARS-CoV-2 ORF1ab (green triangle), N-gene (red square), and S-gene (blue circle), and MS2 (open circle). Undetermined Ct values are plotted at 0.

**Supporting Figure 8.**
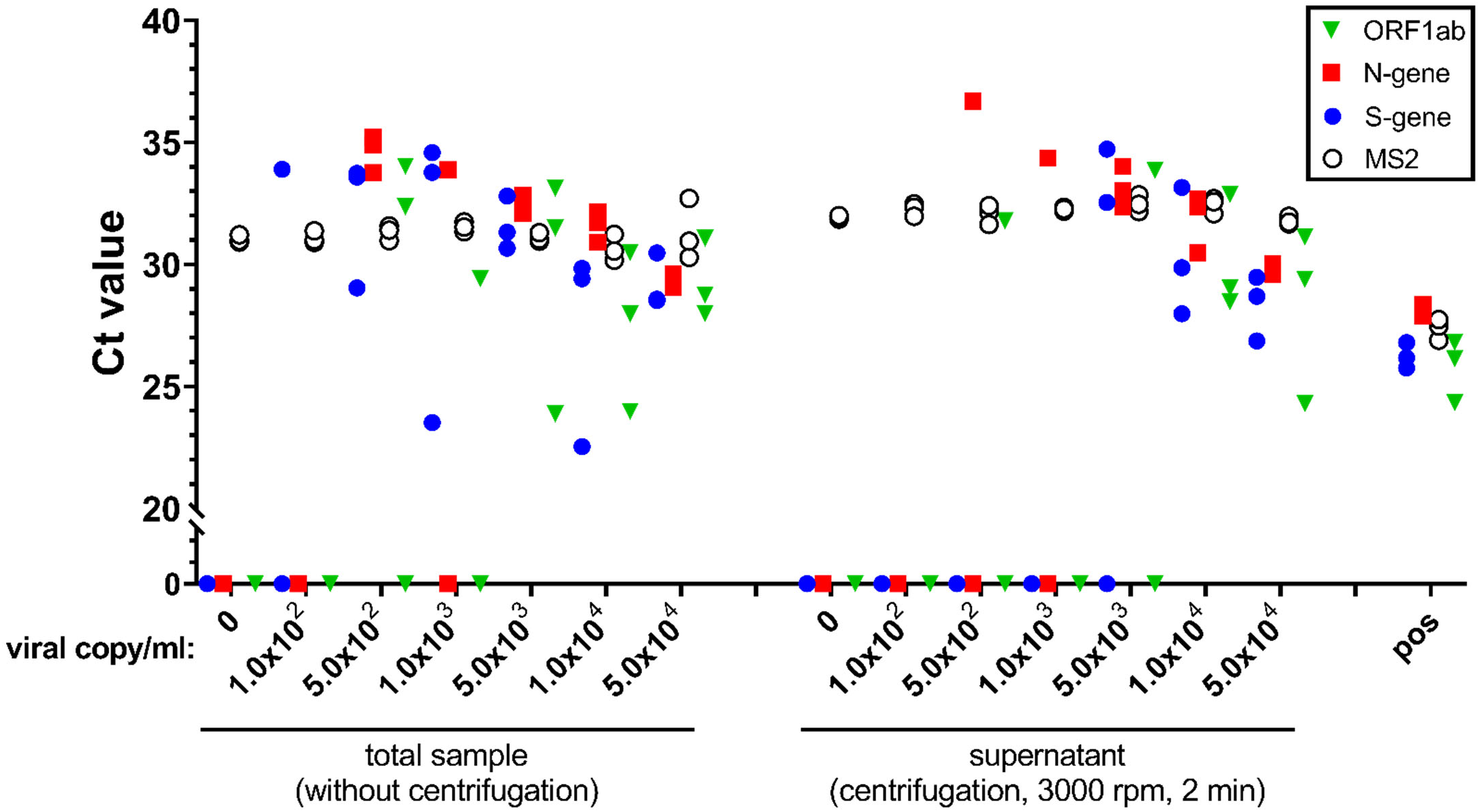
Effect of centrifugation on SARS-CoV-2 detection. Heat-inactivated SARS-CoV-2 (1.0×10^2^, 5.0×10^2^, 1.0×10^3^, 5.0×10^3^, 1.0×10^4^, and 5.0×10^4^ viral copies/mL) was spiked into fresh human saliva (SARS-CoV-2 negative) and combined with TBE buffer 1:1 at a final working concentration of 1X. Samples were heat treated at 95°C for 30 min, then treated with or without centrifugation at 3000 rpm for 2 min. All saliva samples were spiked with purified MS2 bacteriophage (1:40 MS2:sample) as an internal control. Virus-spiked saliva samples, centrifugation supernatants, a positive control (pos; SARS-CoV-2 positive control, 5.0×10^3^ copies/mL) and a negative control (neg; water) were directly analyzed by RT-qPCR, in triplicate, for SARS-CoV-2 ORF1ab (green triangle), N-gene (red square), and S-gene (blue circle), and MS2 (open circle). Undetermined Ct values are plotted at 0.

**Supporting Figure 9.**
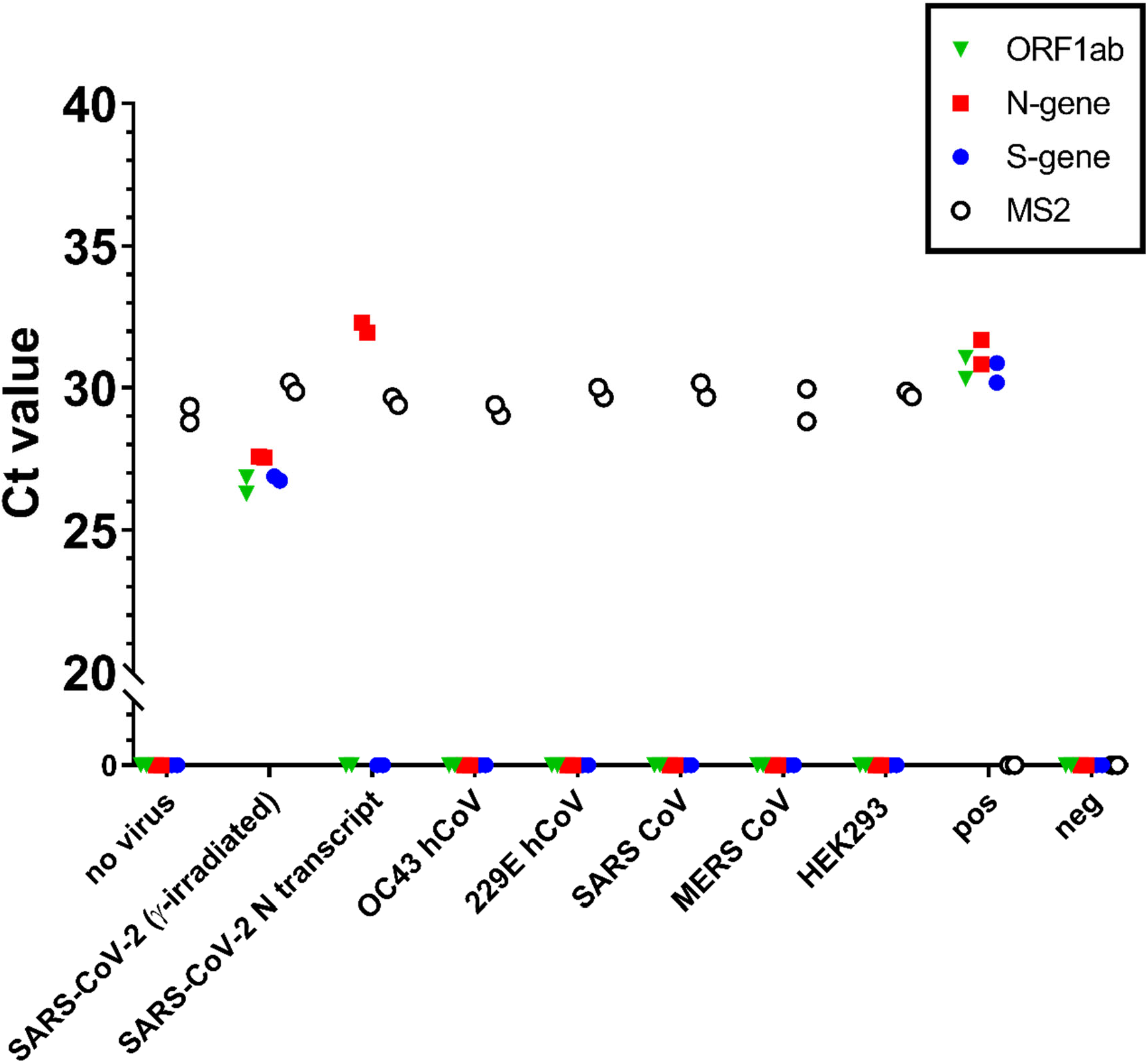
Specificity of SARS-CoV-2 detection system. Commercially available saliva (Lee Biosciences and Innovative Research) were combined in equal proportions, diluted 1:1 with 2X TBE buffer, and spiked 1.0×10^5^ viral copies/mL of SARS-CoV-2 (γ-irradiated virus or synthetic N-transcript RNA), human coronaviruses (229E, OC43), SARS and MERS synthetic RNA, and human RNA (purified from HEK 293 cells). Samples were heat treated at 95°C for 30 min. All saliva samples were spiked with purified MS2 bacteriophage (1:40 MS2:sample) as an internal control. Virus-spiked saliva samples, a positive control (pos; SARS-CoV-2 positive control, 5.0×10^3^ copies/mL, no MS2) and a negative control (neg; water, no MS2) were directly analyzed by RT-qPCR, in triplicate, for SARS-CoV-2 ORF1ab (green triangle), N-gene (red square), and S-gene (blue circle), and MS2 (open circle). Undetermined Ct values are plotted at 0.

